# Signal transduction pathways controlling *Ins2* gene activity and beta cell state transitions

**DOI:** 10.1101/2024.06.06.597838

**Authors:** Chieh Min Jamie Chu, Bhavya Sabbineni, Haoning Howard Cen, Xiaoke Hu, WenQing Grace Sun, George P. Brownrigg, Yi Han Xia, Jason Rogalski, James D. Johnson

## Abstract

Pancreatic β cells exist in low and high insulin gene activity states that are dynamic on a scale of hours to days. Here, we used live 3D imaging, mass spectrometry proteomics, and targeted perturbations of β cell signaling to comprehensively investigate *Ins2*(GFP)^HIGH^ and *Ins2*(GFP)^LOW^ β cell states. We identified the two *Ins2* gene activity states in intact isolated islets and showed that cells in the same state were more likely to be nearer to each other. We report the proteomes of pure β cells to a depth of 5555 proteins and show that β cells with high *Ins2* gene activity had reduced β cell immaturity factors, as well as increased translation. We identified activators of cAMP signaling (GLP1, IBMX) as powerful drivers of transitions from *Ins2*(GFP)^LOW^ to the *Ins2*(GFP)^HIGH^ states. Okadaic acid and cyclosporine A had the opposite effects. This study provides new insight into the proteomic profiles and regulation of β cell states.

## Introduction

Diabetes is characterized by dysfunctional glucose homeostasis which is caused by a relative (type 2 diabetes) or near complete (∼80%; type 1 diabetes) deficiency in insulin, the body’s main anabolic hormone^1^. Insulin is produced by pancreatic islet β cells, whose dysfunction and/or death are prerequisites for diabetes progression^2–4^. Not all β cells are the same^5–7^. Several studies have shown that insulin content and mRNA can vary greatly between individual β cells^8,9^. Our previous live cell imaging using a dual *Ins1* and *Pdx1* fluorescent promoter reporter showed that mouse and human β cell lines could transition from less to more mature states^10,11^ and identified endogenous and chemical regulators of this process^12,13^. However, reporter constructs using abbreviated promoter fragments may not accurately report endogenous gene activity. Therefore, we also used *Ins*2^GFP^ knock-in mice^14^ to investigate endogenous *Ins2* gene activity and identified two distinct β cell states, one with high insulin gene activity and higher insulin mRNA, *Ins2*(GFP)^HIGH^, and one with lower insulin gene activity and lower insulin mRNA, *Ins2*(GFP)^LOW^ ^15^. Live cell imaging confirmed that β cells had the capacity to transition between the high and low GFP states in both directions^15^. Single cell RNA sequencing (scRNAseq) demonstrated *Ins2*(GFP)^HIGH^ cells had a more mature β cell transcriptome profile but were also more fragile^15^. These results suggest that the *Ins2*(GFP)^HIGH^ cells produce more insulin but are also more vulnerable to insults. However, it has been shown that the transcriptome and proteome have weak correlation in islets^16^, so it is critical to understand the proteomic profile of these states. It is also important to understand the molecular mechanisms that control β cell state transitions. ER stress and other cellular perturbations are also known to impact gene transcription^17,18^. For example, stresses associated with the high *Ins2* state lead to β cell fragility^15^ and lower proliferation capacity^19^, but the effects of stress itself on *Ins2* gene activity remains unclear. With robotic imaging technology and automated analysis, it is possible to study dozens of signaling pathways at once.

Live imaging of β cell states presented the opportunity for fluorescence activated cell sorting (FACS) purification of selected cell populations, while advances in mass spectroscopy permit the identification and cataloguing of thousands of cell-selective proteins that could be effects markers or targets for drug delivery. Previous proteomic studies purified β cells using insulin immunofluorescence staining or FAD autofluorescence^20,21^, but these methods for enriching β cells have caveats such as the reliance on antibody specificity and that FAD autofluorescence may not identify all β cells. Thus, there was also a need to revisit those approaches with more sensitive proteomics equipment and analysis pipelines.

Here, we define the proteomic profiles of purified rodent β cells and of the two *Ins2* gene activity states using *Ins2*^GFP^ knock-in mice. We also systematically tested the effects of 48 drugs known to affect selected signaling pathways and cellular functions on *Ins2* gene activity and β cell state transitions. We present a comprehensive comparison of the two *Ins2* expression states at the protein and functional level, and reveal multiple genes implicated in mechanisms related to *Ins2* gene activity and β cell state transitions.

## Results

### Live 3D imaging of intact islets from Ins2^GFP^ mice

Our previous analyses of *Ins2* gene activity in dispersed mouse islets revealed a bimodal distribution of GFP fluorescence from a reporter construct knocked into the endogenous *Ins2* gene locus^15^ (Fig. 1A, B). In this study, we used live 3D imaging to define the prevalence and spatial relationship of cells in the two β cell insulin gene activity states in whole intact islets from *Ins2*^GFP^ knock-in mice (Fig. 1C). We analyzed islets which had below 400 GFP positive cells, as we wanted to avoid large islets which may have necrotic cores^22^. Consistent with dispersed cells, *Ins2* gene activity in intact islets also had a bimodal distribution (Fig. 1D). We also found that the range of GFP fluorescence was greater in larger islets compared to smaller ones (Fig. 1D). We did not find significant sex differences when comparing the ratios of *Ins2*(GFP)^HIGH^ and *Ins2*(GFP)^LOW^ cells in intact male and female islets (Fig. 1E), consistent with our previous work in dispersed β cells^23^. Nearest neighbor analysis showed that cells in the same *Ins2* activity state were more likely to be found in proximity of each other (Fig. 1F; Supplemental Fig. 1A). We found no significant difference between the cell states and their distance to the core of the islet (Fig. 1G; Supplemental Fig. 1B). To summarize, live cell 3D imaging analysis revealed the that the *Ins2*(GFP)^HIGH^ and *Ins2*(GFP)^LOW^ expression states exist in intact islets, with cells in the same activity states being more likely to be found next to each other.

**Figure 1.**
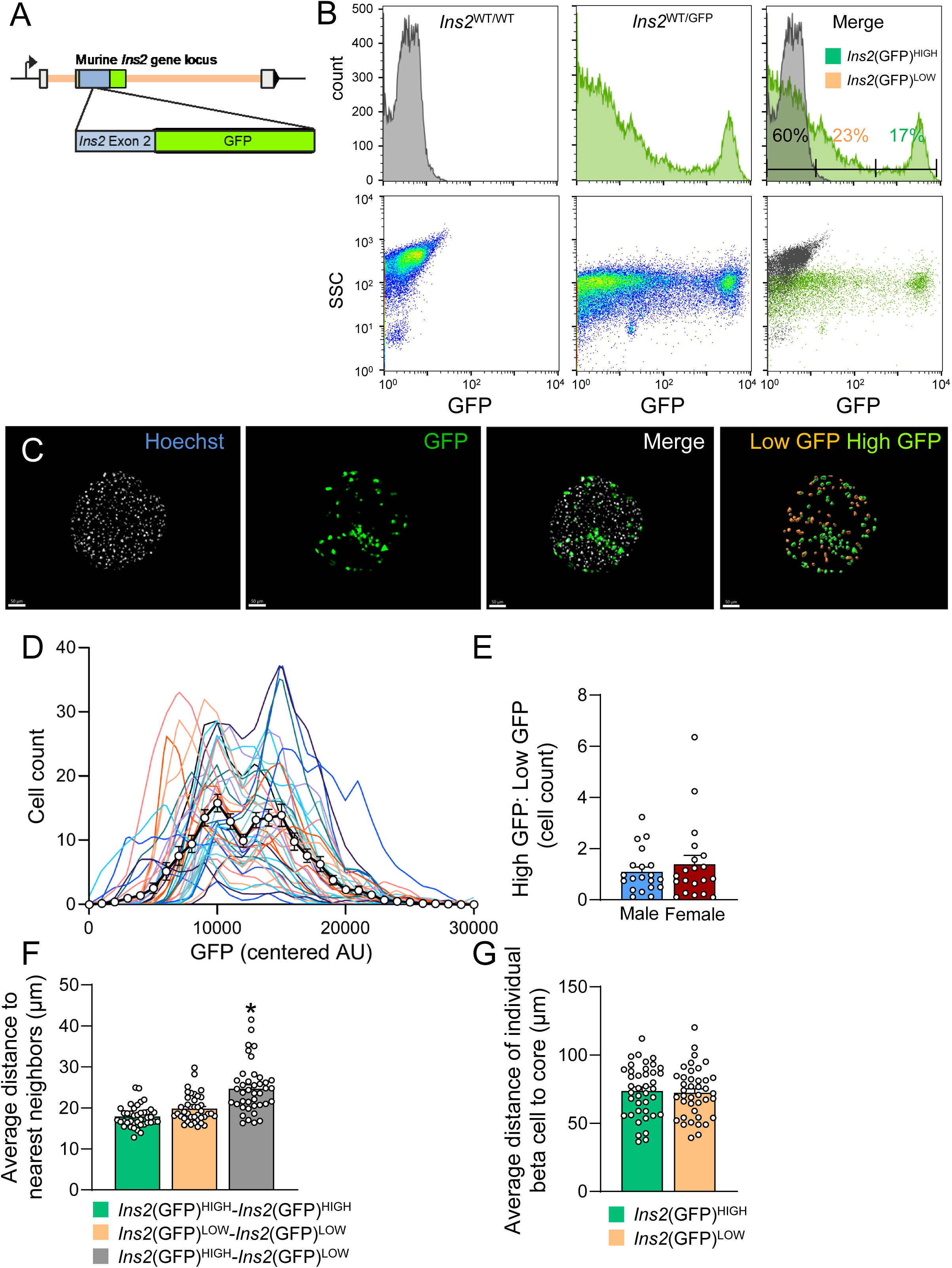
Live 3D imaging of intact *Ins2*^GFP^ mouse islets. **(A)** Schematic of the *Ins2*^GFP^ knock-in mouse model, in which the second exon of the wildtype *Ins2* gene has been partially replaced with GFP. **(B)** FACS purification of *Ins2*(GFP)^LOW^ and *Ins2*(GFP)^HIGH^ populations of β cells, with β cells isolated from a WT mouse for comparison. **(C)** Example images of 3D live cell imaging of intact islets from *Ins2*^GFP^ knock-in mice. Hoechst is a cell nuclei dye. Scale bar is 30 μm. **(D)** Distribution of cells based on GFP fluorescence from *Ins2*^GFP^ islets. Each distribution line represents an islet. (n = 40 islets). **(E)** Comparison of ratios of high GFP cells to low GFP cells in male and female mice. Each dot represents an islet (n = 20 male and 20 female islets from 3 male and 3 female mice). **(F)** Nearest neighbor analysis of cells from *Ins2*^GFP^ knock-in mice (n = 40 islets). One-way ANOVA. **(G)** Distance to core of cells from *Ins2*^GFP^ knock-in mice. Student’s T-test. Density: probability density function, represents the probability a data is located within the defined range (i.e. ratio of the total). Data are represented as mean +/- SEM. * p < 0.05

### Identification of β cell selective islet proteins

Proteins that are selective to β cells have potential as biomarkers and anchor targets for imaging and drug delivery^24^. Previous studies have identified β cell-specific mRNAs using single cell RNA sequencing^25^. However, mRNA and proteins in isolated islets only have a correlation coefficient of ∼0.5^16^, so there is an unmet need for minimally biased, high-throughput characterization of β cell selective gene products at the protein level. Therefore, as a step towards identifying β cell specific proteins, we compared the proteomes of GFP positive cells to GFP negative cells from dispersed and sorted mouse islets to a depth of 5555 proteins (Supplemental Fig. 2A, 3A). When adjusted for multiple comparisons, 20 proteins were found to be significantly upregulated in GFP negative cells (Fig. 2A). These include brain abundant membrane attached signal protein 1 (BASP1), a protein that was found to be upregulated in the islets of non-pregnant mice when compared with pregnant^26^; calbindin 1 (CALB1), gene that was enriched in δ and γ cells^27^; clusterin (CLU), a protein known to play a regenerative role after islet injury^28^; and glutathione peroxidase 3 (GPX3), a regulator of oxidation that may play a role in T2D associated stress (Fig. 2A)^29^.

**Figure 2.**
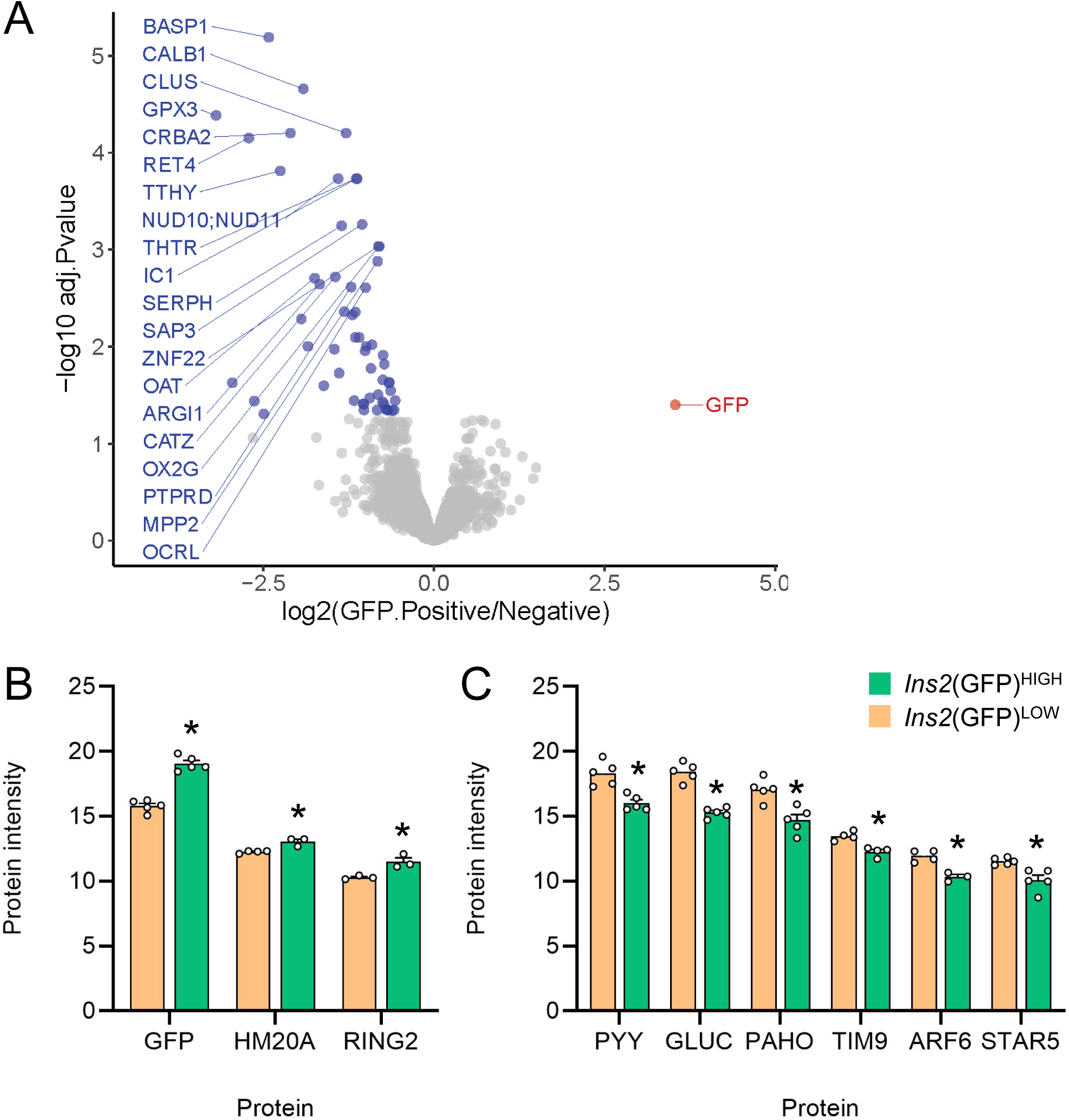
Proteomic analysis of FACS purified *Ins2*^GFP^ β cells. **(A)** Volcano plot of proteins differentially abundant when comparing GFP positive cells with GFP negative cells. **(B)** Barplot of proteins differentially upregulated in *Ins2*(GFP)^HIGH^ cells. **(C)** Barplot of proteins differentially upregulated in *Ins2*(GFP)^LOW^ cells. FACS purified samples were acquired from 3 male and 2 female mice.

We also undertook a more liberal analysis that did not account for multiple comparisons on our proteomics data. As expected, GFP-negative, non-β cells were enriched for known markers of other islet cell types, such as GCG, PYY, and PPY (Supplemental Fig. 4A). To rank the proteins, we multiplied the p-value by the fold change. The top novel non-β cell proteins were the cell surface protein tetraspanin 8 (TSPAN8), arginase 1 (ARG1), and clusterin (CLU) (Fig. 2A). The top 5 most significantly enriched β cell selective proteins were: 1) CAP-Gly domain containing linker protein 1 (CLIP1), which links cytoplasmic vesicles to microtubules; 2) 3’-phosphoadenosine 5’- phosphosulfate synthase 2 (PAPSS2), which mediates two steps in the sulfate activation pathway; 3) tyrosine 3-monooxygenase/tryptophan 5-monooxygenase activation protein zeta (YWHAZ), also known as 14-3-3ζ, a scaffolding protein we have shown plays important survival and signaling roles in β cells^30^; 4) apoptosis inducing factor mitochondria associated 1 (AIFM1); and 5) cystathionine beta synthase (CBS), a gene which may play a role in β cell insulin secretion^31^ (Supplemental Fig. 4A). As expected, INS1 and IAPP proteins were also significantly enriched in pure β cells. The G protein-coupled receptor 158 (GPR158), a metabotropic glycine receptor^32^ which may play a role in autocrine feedback^33^, was highly enriched in β cells.

We next undertook pathway analysis to identify themes in our proteomic data from non-β cells and pure β cells (Supplemental Fig. 4B). Gene set enrichment analyses revealed that non-β cells are enriched for proteins that mediate metabolic and catabolic processes including ‘cellular modified amino acid metabolic process’ and ‘carbohydrate derivative metabolic process’. In pure β cells, there was enrichment of pathways entitled ‘skin development’ and ‘supramolecular fiber organization’, which was driven by the relative abundance of multiple keratin family proteins, which are known to modulate β cell signalling and play roles in stress responses^34^. Enrichment of ‘regulation of cellular component biogenesis’ and ‘regulation of protein-containing complex assembly’ pathways suggests β cells may exist in a more anabolic state than non-β cells. Pathways involving ‘protein phosphorylation’ and ‘IκB/NFκB signalling’ were also enriched in pure GFP-positive β cells (Supplemental Fig. 4B). Interestingly, NFκB signalling has been implicated in a β cell subpopulation with lower PDX1 expression, which has similarities with our *Ins2*(GFP)^LOW^ cells^35^. Overall, these results suggest that differences between *Ins2* expressing cells and other islet cells are primarily associated with protein production and signalling.

### Proteomic profiles of FACS purified β cells in Ins2(GFP)^HIGH^ and Ins2(GFP)^LOW^ states

We next compared the proteomic profiles of the two distinct *Ins2* gene activity states in FACS purified β cells (Supplemental Fig. 5A). After adjusting for multiple comparisons, we found that, other than GFP, two proteins were significantly upregulated in *Ins2*(GFP)^HIGH^ cells: HMG20A, a protein associated with type 2 diabetes and shown to be important to functional maturity of β cells^36^; and ring finger protein 2 (RING2), a member of the polycomb group of proteins which are important for transcriptional repression of genes involved in cell proliferation and development^37^. In the *Ins2*(GFP)^LOW^ state, proteins with greater abundance include the β cell immaturity markers pancreatic polypeptide (PPY), peptide YY (PYY), and glucagon (GCG)^38^; translocase Of inner mitochondrial membrane 9 (TIM9), a protein that helps mediate mitochondrial protein import^39^; ADP ribosylation factor 6 (ARF6), a protein that has been shown to mediate glucose stimulated insulin secretion^40^; and stAR-related lipid transfer domain containing 5 (STARD5), a ER stress protein which helps regulate cholesterol homeostasis^41^. Overall, these findings further support our hypothesis that cells in the *Ins2*(GFP)^HIGH^ state have a more mature β cell profile.

We then analyzed our data without adjusting for multiple comparisons to get an idea of the trends of proteomes of the two cell states. Using p value times fold change as a criteria again, the top 5 most enriched proteins in *Ins2*(GFP)^LOW^ cells were: 1) glucagon (GCG), a marker for alpha cells; 2) pancreatic polypeptide (PPY) a marker for PP cells; 3) Purkinje cell protein 4 (PCP4), a calmodulin regulator involved in calcium binding and signalling; 4) aldehyde dehydrogenase 1 family member 3 (ALDH1L2), a mitochondrial enzyme implicated in β cell protection and secretory dysfunction ^42^; and 5) adenine phosphoribosyltransferase (APRT), an enzyme involved in the nucleotide salvage pathway (Supplemental Fig. 5A). These results suggest that cells in the less mature *Ins2*(GFP)^LOW^ state was enriched in markers of ‘multi-hormonal’ cells, consistent with our previous scRNAseq analysis ^15^. We also found enrichment in proteins in endoplasmic reticulum processing, including HSPA5, HSP90B1, and ERO1L (Supplemental Fig. 8A-C), corroborating our prior scRNAseq analyses^15^.The top 5 most enriched proteins (other than GFP) in *Ins2*(GFP)^HIGH^ cells were: 1) zinc finger AN1-type containing 2B (ZFAND2B), a protein associated with the proteasome for degradation of misfolded proteins; 2) zinc finger RANBP2-type containing 2 (ZRANB2), a protein involved in RNA splicing and mRNA processing 3) myocyte enhancer factor 2D (MEF2D), a calcium regulated transcription factor; 4) SWI/SNF related, matrix associated, actin dependent regulator Of chromatin subfamily A, member 2 (SMARCA2), a protein which regulates transcription via altering chromatin structure; and 5) phosphofurin acidic cluster sorting protein 2 (PACS2), a protein involved in ER calcium ion homeostasis. The protein poly(A) polymerase alpha (PAPOLA), a protein required for creation of the 3’-poly(A) tail of mRNAs, was also upregulated in *Ins2*(GFP)^HIGH^ cells (Supplemental Fig. 5A).

The pathways significantly enriched in the *Ins2*(GFP)^LOW^ state relative to the *Ins2*(GFP)^HIGH^ state included: ‘organic acid metabolic process’, ‘cellular respiration’, ‘generation of precursor metabolites and energy’, and ‘oxidative phosphorylation’, suggesting the less mature cells may have more energic capacity. They are also enriched in proteins involved in ‘fatty acid metabolic processes’ and ‘lipid metabolic processes’, suggesting the possibility of metabolic substrate preference and switching between these cell states (Supplemental Fig. 5B). The *Ins2*(GFP)^LOW^ state was enriched in ‘hormone metabolic process’ which was also enriched in non-β cells relative to β cells, including GCG, PYY, and SST. Gene set enrichment analysis revealed that the pathways most enriched in the *Ins2*(GFP)^HIGH^ state were related to transcriptional upregulation and mRNA processing (Supplemental Fig. 5B). The upregulation of ‘DNA-templated transcription’ and ‘RNA biosynthetic process’ pathways may suggest that the *Ins2*(GFP)^HIGH^ state is more transcriptionally active. The *Ins2*(GFP)^HIGH^ state also enriched for ‘RNA processing’, a pathway which involves mRNA capping and addition of the 3’poly(A) tail. Regulation and processing of the 5’ and 3’ UTRs of insulin mRNA is known for their importance in maintaining mRNA stability, and our prior scRNAseq and qPCR data indeed showed that the *Ins2*(GFP)^HIGH^ state had more *Ins2* mRNA^15^. ‘Chromosome organization’ and ‘cell cycle’ pathway upregulation is also indicative of regulation of the transcriptional landscape. Overall, this data suggests that differences between the *Ins2*(GFP)^HIGH^ and *Ins2*(GFP)^LOW^ states were related to transcription mechanisms and processes.

We next analysed the relationship between our differentially abundant proteins using STRING and mapped them to a cell diagram in Cytoscape to visualize protein-protein networks in their most common cellular locations (Supplemental Fig. 6A, B). Protein-protein interaction networks enriched in GFP negative cells, beyond the known markers from other islet cell types (GCG, PYY, PPY, SST), included protein networks related to lipid metabolism (GPX3, HEXB) and factors related to amino acid synthesis and transport (SLC7A8, SLC3A2, ARG1) (Supplemental Fig. 6A). Protein-protein interaction networks enriched in GFP positive cells, compared with non-GFP, included those involved in GPCR and cAMP signalling, glycolysis, and the secretory pathway (ER, Golgi, granules, cytoskeleton). Specifically, pure β cells showed enrichment for multiple cytoskeletal factors, including VIL1, a member of calcium regulated actin binding proteins and is known plays a role in insulin exocytosis regulation; and CLIP1, a protein associated with granule trafficking^43–45^. These data provide data on β cell specific protein networks at unprecedented detail.

Next, we examined network differences between the *Ins2*(GFP)^HIGH^ state and the *Ins2*(GFP)^LOW^ state. The *Ins2*(GFP)^LOW^ state was enriched for networks related to mitochondria function and cell respiration, including members of the pyrroline-5-carboxylate reductase family (PYCR1, PYCR2), and subunits of the NADH dehydrogenase (ubiquinone) complex (NDUFA6, NDUFB6, NDUFB10) (Supplemental Fig. 7A). We also noted several factors with links to proteasome function (PSMA3, PSMB3, PSMB8), and translation factors (EIF3BEIF4E, EIF4A1) (Supplemental Fig. 7A). This suggests that cells with low *Ins2* gene activity may be recovering from various sources of stress and may also be in the process of regulating translation machinery. Interestingly, RHOA, an F-actin regulator which reduces insulin secretion when activated, was upregulated in the *Ins2*(GFP)^LOW^ state ^46^. Another actin protein, ARPC2, which was downregulated in human islets when treated with insulin, was also enriched in *Ins2*(GFP)^LOW^ cells^47^. On the other hand, many of the enriched proteins in the *Ins2*(GFP)^HIGH^ state were located in the nucleus and were linked to transcriptional regulation, including prominent transcription factors TOP1, CDK9, and KAT7(Supplemental Fig. 7B). Several nuclear proteins were associated with mRNA processing, such as ZRANB2, SRPK1, and PAPOLA^48–50^ (Supplemental Fig. 7B). We also noted a number of factors in the Golgi, including STRIP1, a protein involved in the localization of the Golgi, and VCPIP1, a protein involved in Golgi reassembly. There were also a fair number of proteins in the ER, including the aforementioned ZFAND2B and PACS2; DNAJA1, a protein which stimulates ATP hydrolysis; and NGLY1, an enzyme involved in the degradation of misfolded proteins. Taken together, the *Ins2*(GFP)^HIGH^ state is primarily associated with the regulation of transcription, mRNA processing, and protein trafficking, while the *Ins2*(GFP)^LOW^ state may be in the process of recovering from stress associated with high *Ins2* gene activity and a shift in protein translation machinery.

Our proteomics data revealed enrichment of ‘regulation of cellular component biogenesis’ and ‘regulation of protein-containing complex assembly’ pathways in the *Ins2*(GFP)^HIGH^ state, so we tested whether β cell translation capacity was different compared to other islet cell types using the O-propargyl-puromycin (OPP) assay, which incorporates a fluorescent marker into nascent proteins (Fig. 3A). Using GFP as a marker for β cells, we found that β cells had higher translational capacity compared to other islet cells (Fig. 3B). The upregulation of DNA template transcriptional regulation and RNA processing pathways in *Ins2*(GFP)^HIGH^ cells relative to *Ins2*(GFP)^LOW^ cells is suggestive of increased downstream translational capacity. We found that cells in the *Ins2*(GFP)^HIGH^ state had significantly higher translational capacity compared to cells in the *Ins2*(GFP)^LOW^ state (Fig. 3C, D). We did not note any significant differences when comparing the translational capacity of the two *Ins2* expression states in the context of sex differences (Supplemental Fig. 10A). Overall, these data reveal the differences between the *Ins2*(GFP)^HIGH^ and *Ins2*(GFP)^LOW^ expression states at the protein level, and show how some of these differences can translate to differences in β cell function.

**Figure 3.**
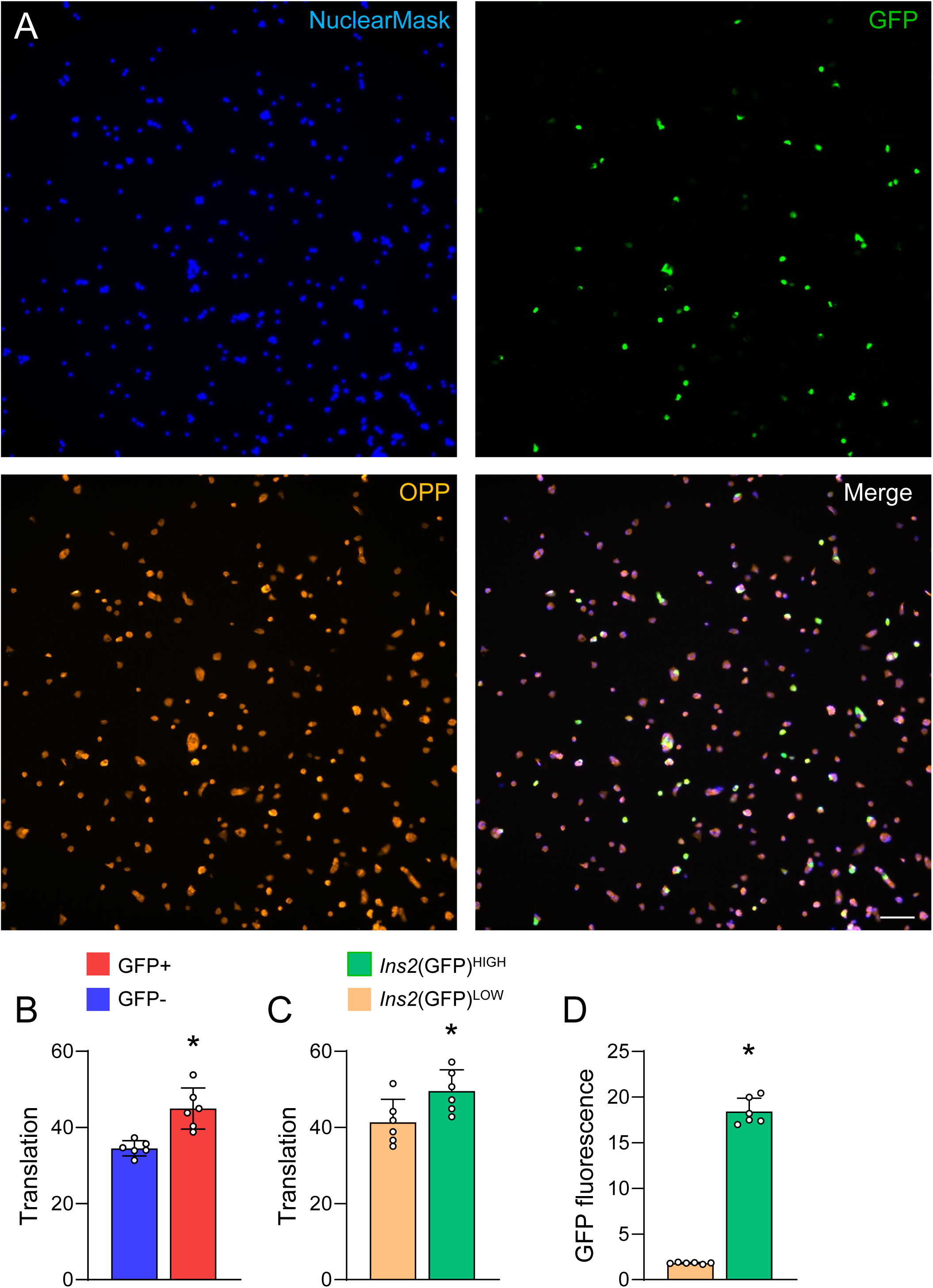
OPP protein synthesis assay of *Ins2*(GFP)^LOW^ and *Ins2*(GFP)^HIGH^ cells. **(A)** Representative images of OPP protein synthesis assay on *Ins2*^WT/GFP^ mice. The blue wavelength is the Nuclearmask cell nuclei stain, the green wavelength is GFP, and the red wavelength is the OPP marker for nascent peptide formation. Scale bar is 40 μm. **(B)** Comparison of translational capacity of GFP positive β cells and GFP negative cells (n = 6 mice, 3 males and 3 females). Student’s t-test. **(C)** Comparison of translational capacity of *Ins2*(GFP)^HIGH^ and *Ins2*(GFP)^LOW^ cells (n = 6 mice, 3 males and 3 females). Student’s t-test. **(D)** GFP fluorescence of *Ins2*(GFP)^LOW^ β cells and *Ins2*(GFP)^HIGH^ β cells. Student’s t-test (n = 6 mice, 3 males and 3 females). Data are represented as mean +/- SEM. * p < 0.05.

### Mechanistic interrogation of Ins2 gene activity regulation

The molecular mechanisms regulating *Ins2* gene activity remain to be fully elucidated. We chose 48 drugs (Table 1) known to perturb key β cell signaling pathways to study their effects on β cell states (Fig. 4A). Concentrations for the various compounds were decided based on documented doses used in *in vitro* assays. Dispersed islet cells from *Ins2*^GFP^ mice were treated with compounds and imaged in a high-throughput format (Fig. 4B-L). Although all experiments were conducted in parallel using a robotic imaging system, we visualized the results from each family of compounds separately. Compounds which activate cAMP signalling, namely the PDE inhibitor 3-isobutyl-1-methylxanthine (IBMX) and the cAMP inducer GLP-1, significantly increased *Ins2* gene activity when tracked over time (Fig. 4B). The rate of increase for GFP fluorescence was considerably greater in *Ins2*(GFP)^HIGH^ cells in response to cAMP pathway activators compared to *Ins2*(GFP)^LOW^ cells (Fig. 5B, C). Compared to DMSO control, the calcineurin pathway inhibitors cyclosporine A, okadaic acid, and tacrolimus reduced *Ins2* gene activity (Fig. 4C). Inhibitors of the endonuclease IRE1A caused a steep drop in *Ins2* gene activity after prolonged treatment, with APY29 inducing a strong increase prior to the drop (Fig. 4D). Effectors of ER pumps and channels had relatively few effects on *Ins2* gene activity, with ATP2A2 inhibitor thapsigargin slightly increasing *Ins2* activity (Fig. 4E). Compounds which impacted insulin signalling also had some interesting effects, in particular the protein kinase inhibitor staurosporine, which significantly increased *Ins2* gene activity, and PI3K inhibitor LY294002, which caused a spike at 48 hours followed by a steep decline (Fig. 4F). The glucocorticoid agonist dexamethasone reduced *Ins2* activity, while PPARG agonist rosiglitazone slightly increased it (Fig. 4G). The three nutrients we tested all increased *Ins2* gene activity, with arginine and sodium butyrate having a more significant increase over time compared to palmitic acid (Fig. 4H). Compounds which affected cell membrane excitability also had interesting effects, with K_ATP_ channel (ABCC8 subunit) activator diazoxide and CACNA1C inhibitor verapamil decreasing *Ins2* gene activity, while K_ATP_ channel (ABCC8 subunit) inhibitor tolbutamide modestly increased it near the end of our experiment (Fig. 4I). Perturbing protein degradation factors also yielded effects, in particular ERAD inhibitor eeyarestatin I, which decreased *Ins2* gene activity rapidly after treatment (Fig. 4J). As expected, many compounds which affected transcriptional machinery also impacted *Ins2* gene activity (Fig. 4K). TFIIH inhibitor Triptolide and RNA polymerase II inhibitor alpha amanitin resulted in an increase in *Ins2* activity, peaking at around 48 hours and rapidly declining afterwards (Fig. 4K). CDK9 inhibitor 5, 6-dichloro-1-β-D-ribofuranosylbenzimidazole (DRB) also increased *Ins2* gene activity (Fig. 4K). TOP1 inhibitor actinomycin D caused a steep decline in *Ins2* gene activity 24 hours after treatment (Fig. 4K). HDAC inhibitors MGCD0103 and trichostatin A both reduced *Ins2* gene activity, but not as abruptly as triptolide and alpha amanitin (Fig. 4K). Compounds which affected translation that altered *Ins2* gene activity include EEF2 inhibitor cyclohexamide, which increased *Ins2* activity followed by a steep decline after 48 hours, PKR inhibitor, which increased *Ins2* gene activity, and PERK activator CCT020312 and EIF2S1 inhibitor sal003, which reduced *Ins2* gene activity (Fig. 4L).

**Table 1.**
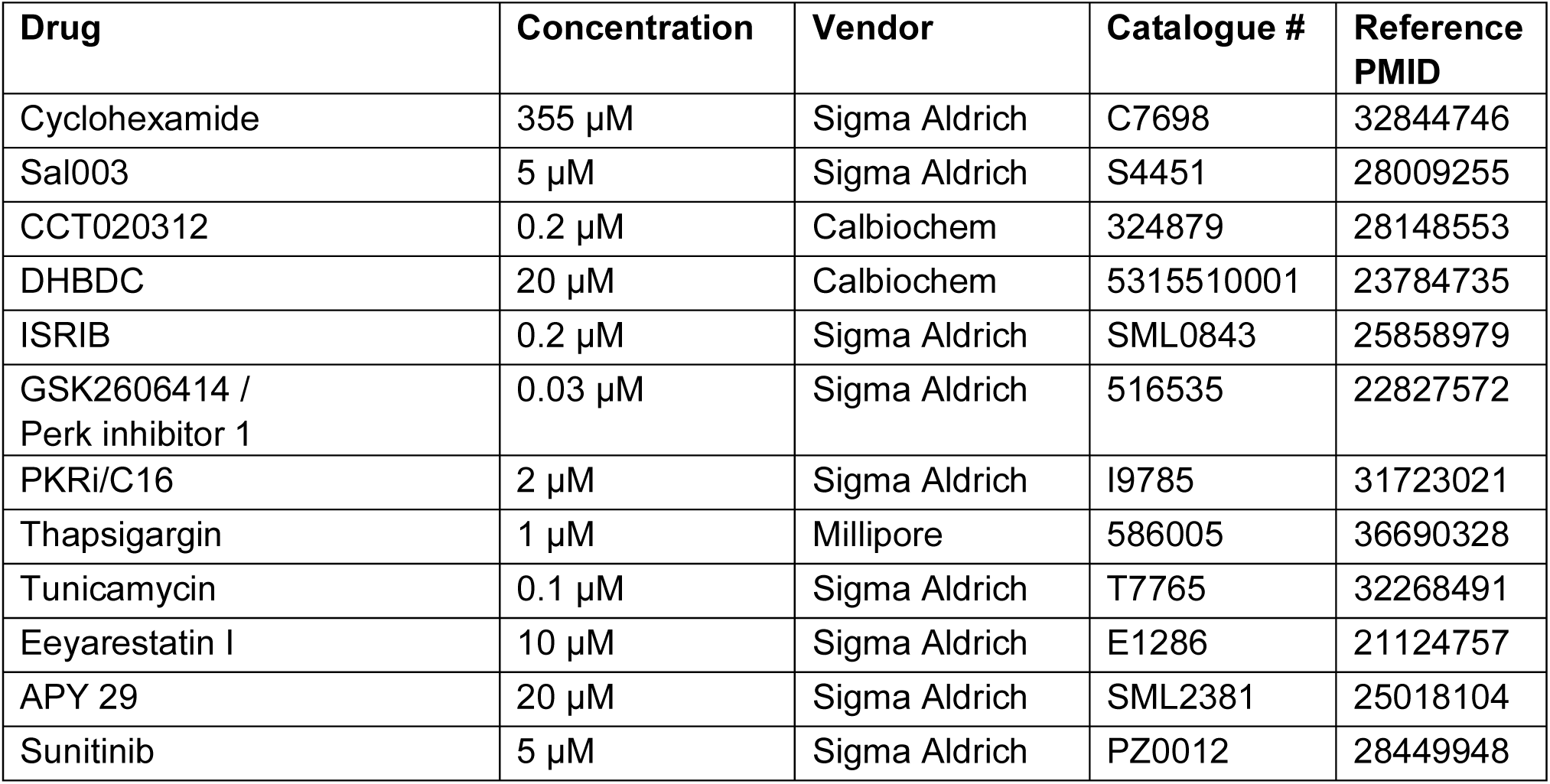
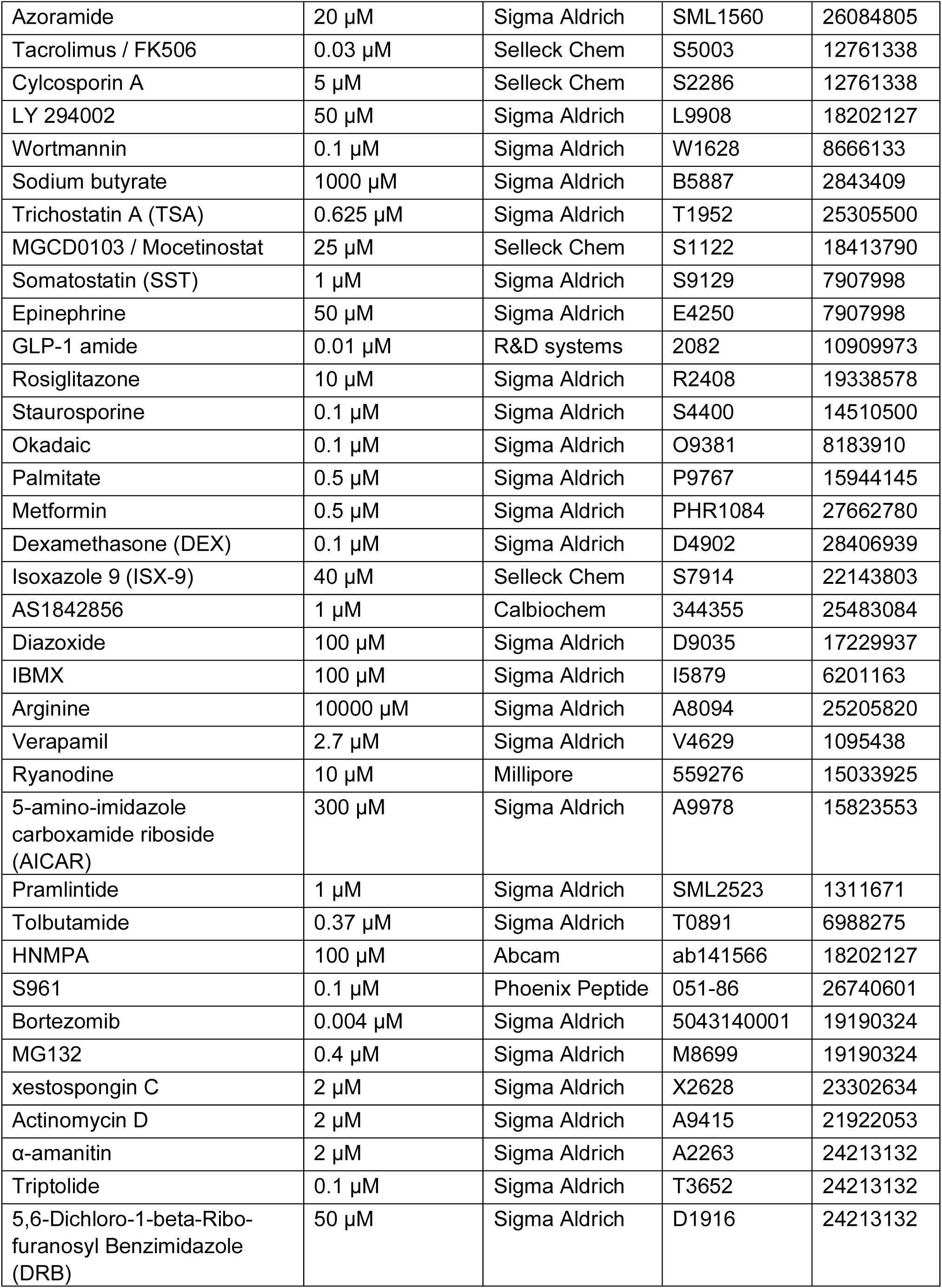
Reagents used in the small molecule screen.

**Figure 4.**
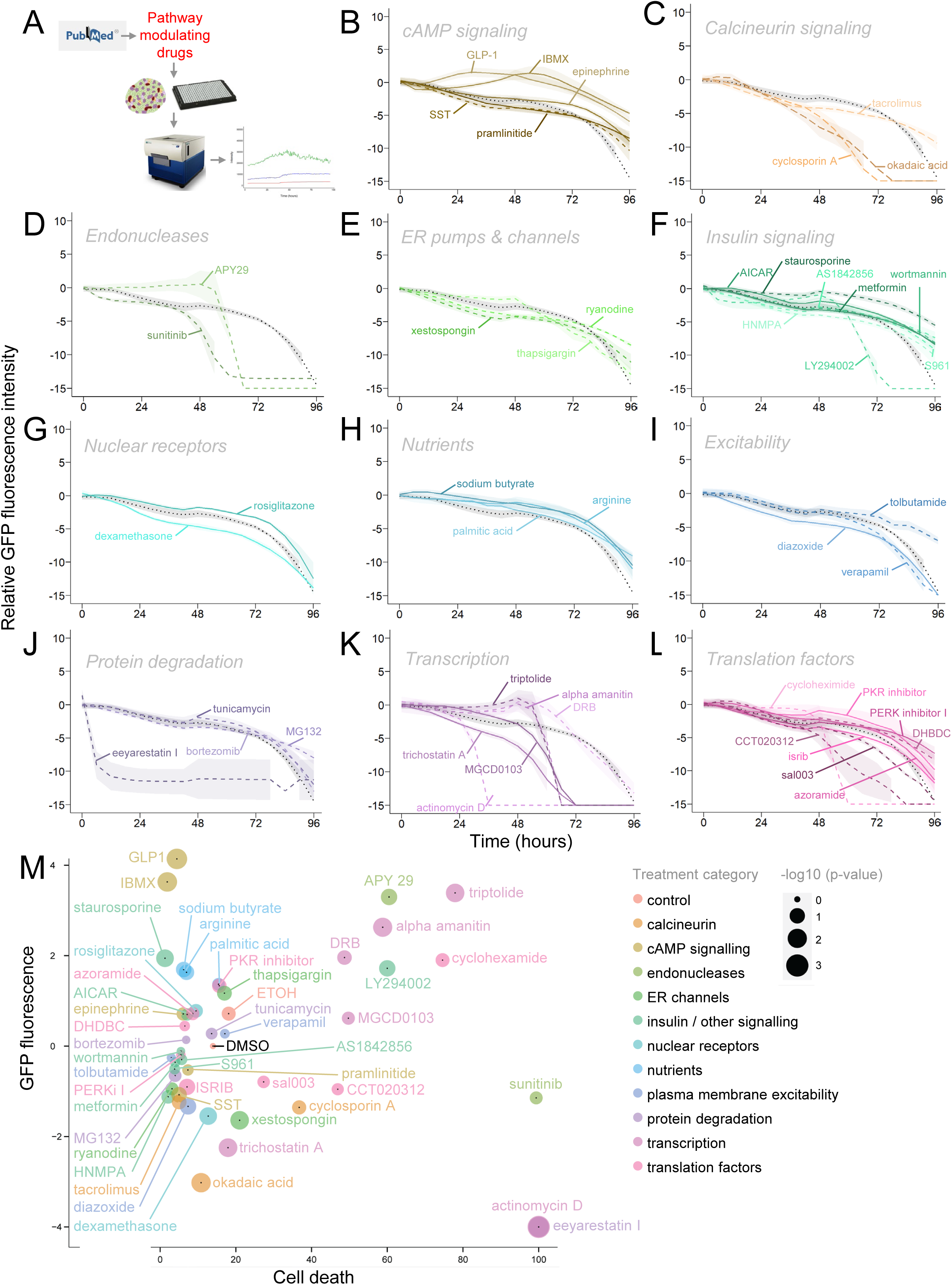
A small molecule screen for effectors of *Ins2* gene activity. **(A)** Experiment pipeline. Islets were isolated and dispersed from *Ins2*^GFP^ knock-in mice. Cells were then stained with Hoechst cell nuclei dye and propidium iodide (PI) cell death marker for two hours, then subjected to treatments. All three fluorescent wavelengths were imaged for 96 hours. **(B-L)** Average GFP fluorescence of cells over the period of 96 hours. Each line represents a treatment. Treatments were categorized based on their primary binding partners. Deceased cells were removed by thresholding for cells with high PI fluorescence. Dotted black lines indicate DMSO controls, dashed lines indicate inhibitors of their respective categories, and solid lines indicate activators of their respective categories (n = 6 mice, 3 males and 3 females). Shading represents SEM. **(M)** Analysis of GFP fluorescence against cell death. GFP fluorescence is scaled GFP fluorescence relative to background after 48 hours of treatment. Cell death is scaled PI fluorescence relative to background after 72 hours of treatment. Size of dots represents p value (adjusted). Colors of dots represents categories, which were decided based on their primary binding partners (n = 6 mice, 3 males and 3 females). Two-way ANOVA, adjusted p value indicated by size of dots.

**Figure 5.**
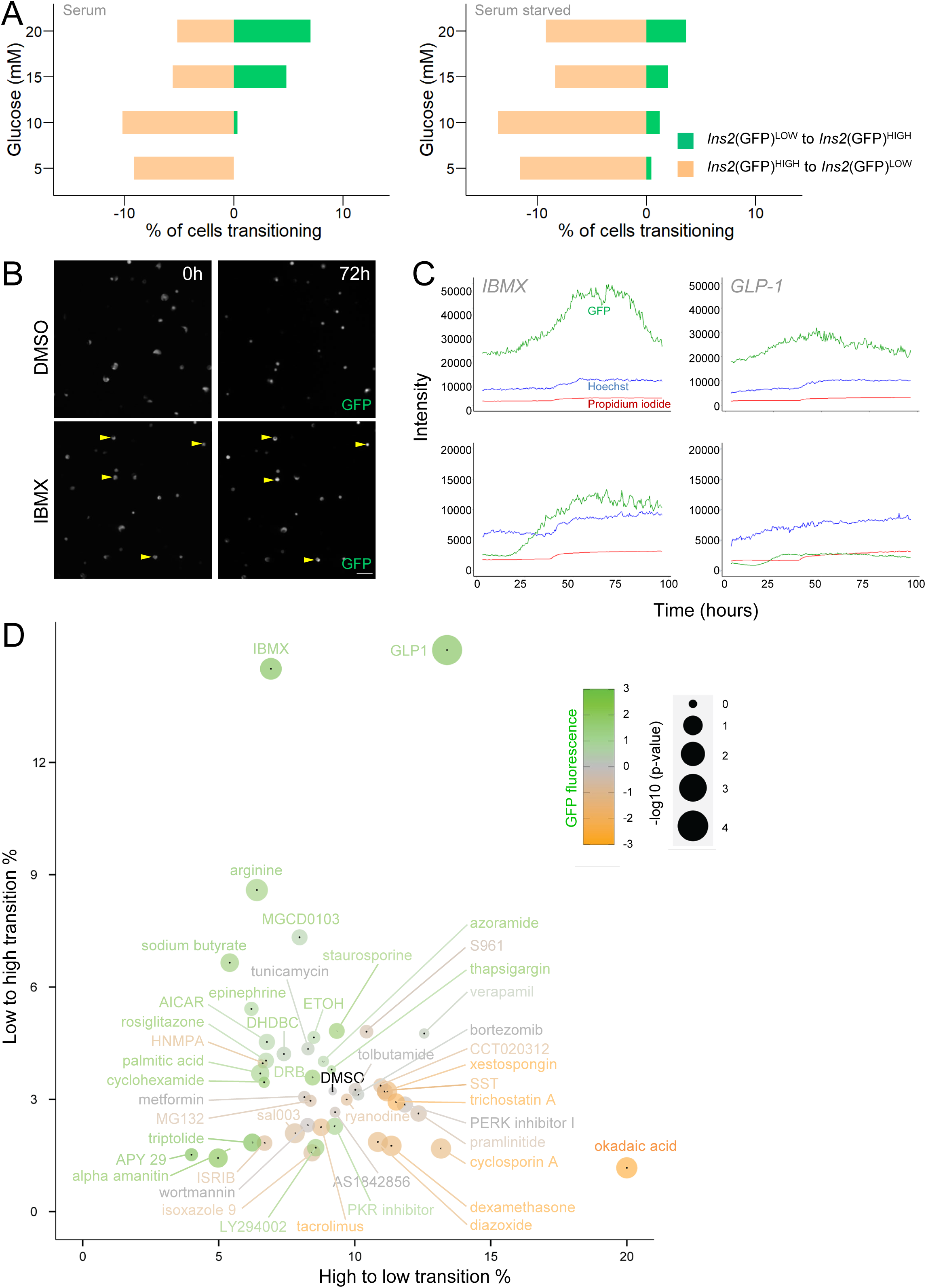
Cell state transitions in the context of various media conditions and perturbations. **(A)** Transition rates between the *Ins2(*GFP)^HIGH^ and *Ins2(*GFP)^LOW^ cell states in the context of various media conditions. % of cells transitioning: percent of subpopulation transitioning to other cell state after 48 hours of culture. **(B)** Example images of increasing GFP fluorescence in response to IBMX. Scale bar is 50 μM. **(C)** Example traces of GFP upregulation and cell state transitions in response to IBMX and GLP-1 amide. **(D)** Comparison of *Ins2(*GFP)^HIGH^ to *Ins2(*GFP)^LOW^ and *Ins2(*GFP)^LOW^ to *Ins2(*GFP)^HIGH^ transition rates in the context of the small molecule screen. Data shown is average transition rate by the end of 48 hours of treatment (n = 6 mice, 3 males and 3 females). Size of the dots denotes p value (adjusted). Color of the dots denotes scaled GFP fluorescence. Two-way ANOVA.

We performed hierarchical clustering using various cell behavior traits in the context of comparing cells from male and female mice. Cell activity traits we used included mean GFP (average GFP fluorescence over time), max GFP (peak GFP fluorescence), and mean cell death (average PI intensity). Overall, the strongest effectors of GFP fluorescence did not show any significant sex differences (Supplemental Fig. 9).

Rapid drops in GFP fluorescence are indicative of cell death, so we explored the relationship between GFP expression and cell viability by plotting fluorescence in the green wavelength (GFP) against the red wavelength (PI) (Fig. 4M)^15^. We note that the cAMP agonists IBMX and GLP-1 amide had the highest GFP fluorescence with relatively low cell death (Fig. 4M). We found that okadaic acid and trichostatin A had the greatest decrease in GFP fluorescence without significant cell death (Fig. 4M). The subset of drugs which had increasing GFP fluorescence followed by a rapid drop (APY 29, triptolide, alpha amanitin, cyclohexamide, DRB, LY294002) had significant cell death, indicating the drugs were toxic after prolonged treatment (Fig. 4M). This is supported by an OPP assay done in the context of several small molecules of interest (GLP-1 amide, IBMX, APY 29, triptolide), in which a significant drop in translational capacity can be observed after 24 hours of treatment, well before the drop in GFP fluorescence and noticeable cell death (Supplemental Fig. 10B). Overall, our small molecule screen shows how various drugs and small molecules may impact *Ins2* gene activity and how it may correlate to β cell health.

### Cell state transitions in the context of media conditions and perturbations

Various culture conditions can impact cell state transitions, with higher glucose concentrations and the addition of FBS inducing a greater rate of *Ins2*(GFP)^LOW^ to *Ins2*(GFP)^HIGH^ transitions (Fig. 5A). We next analyzed the rate of transitions for cells from *Ins2*^GFP^ knock-in mice in the context of our small molecule screen (Fig. 5D). In general, most small molecules had higher *Ins2*(GFP)^HIGH^ to *Ins2*(GFP)^LOW^ transitions rates compared to *Ins2*(GFP)^LOW^ to *Ins2*(GFP)^HIGH^, with our DMSO control at possessing a ∼3:1 ratio (9.1% *Ins2*(GFP)^HIGH^ to *Ins2*(GFP)^LOW^, 3.2% *Ins2*(GFP)^LOW^ to *Ins2*(GFP)^HIGH^). Compared to our DMSO control, we found that cAMP agonists GLP-1 amide and IBMX had the greatest increase in *Ins2*(GFP)^LOW^ to *Ins2*(GFP)^HIGH^ transitions (19% for GLP-1 amide, 14% for IBMX), with relatively low levels of *Ins2*(GFP)^HIGH^ to *Ins2*(GFP)^LOW^ transitions (Fig. 5B-D). Okadaic acid and cycplosporin A induced the highest rate of *Ins2*(GFP)^HIGH^ to *Ins2*(GFP)^LOW^ transitions (Fig. 5D). Interestingly, we found that several drugs with high GFP, including triptolide, APY 29, and alpha amanitin, had very few transitions of either type (Fig. 5D). This shows that the increase in GFP fluorescence observed in these treatments was primarily driven by increasing GFP fluorescence in the *Ins2*(GFP)^HIGH^ cells, and not in the *Ins2*(GFP)^LOW^ cells.

STRING/Cytoscape analyses helped contextualize the major binding partners of our small molecule screen (Fig. 6A, B). Several of the strongest effectors of GFP fluorescence were found to impact genes located within the nucleus, including CDK9, TOP1, POL2A, ERCC3, HDAC1, HDAC2, SP1, and PAK1. Many of these proteins have known roles in transcriptional mechanisms ^51–56^. On the membrane, GLP1R and NOS1 were receptors which had effects on GFP fluorescence. In the ER, perturbing ERN1, a major factor in UPR response, impacted GFP fluorescence and cell health ^57^. In the cytosol, perturbation of VCP, a major component of ERAD, resulted in heavy cell death, while PIK3CA, PDE3A, EEF2 influenced GFP fluorescence. In the context of cell state transitions, perturbing PDE3A and GLP1R induced the highest rate of *Ins2*(GFP)^LOW^ to *Ins2*(GFP)^HIGH^ transitions, while perturbing PPP3CC and PPP2CA resulted in the highest rate of *Ins2*(GFP)^HIGH^ to *Ins2*(GFP)^LOW^ transitions. Overall, these results show how perturbing various β cell signalling pathways may impact *Ins2* gene activity and cell state transitions.

**Figure 6.**
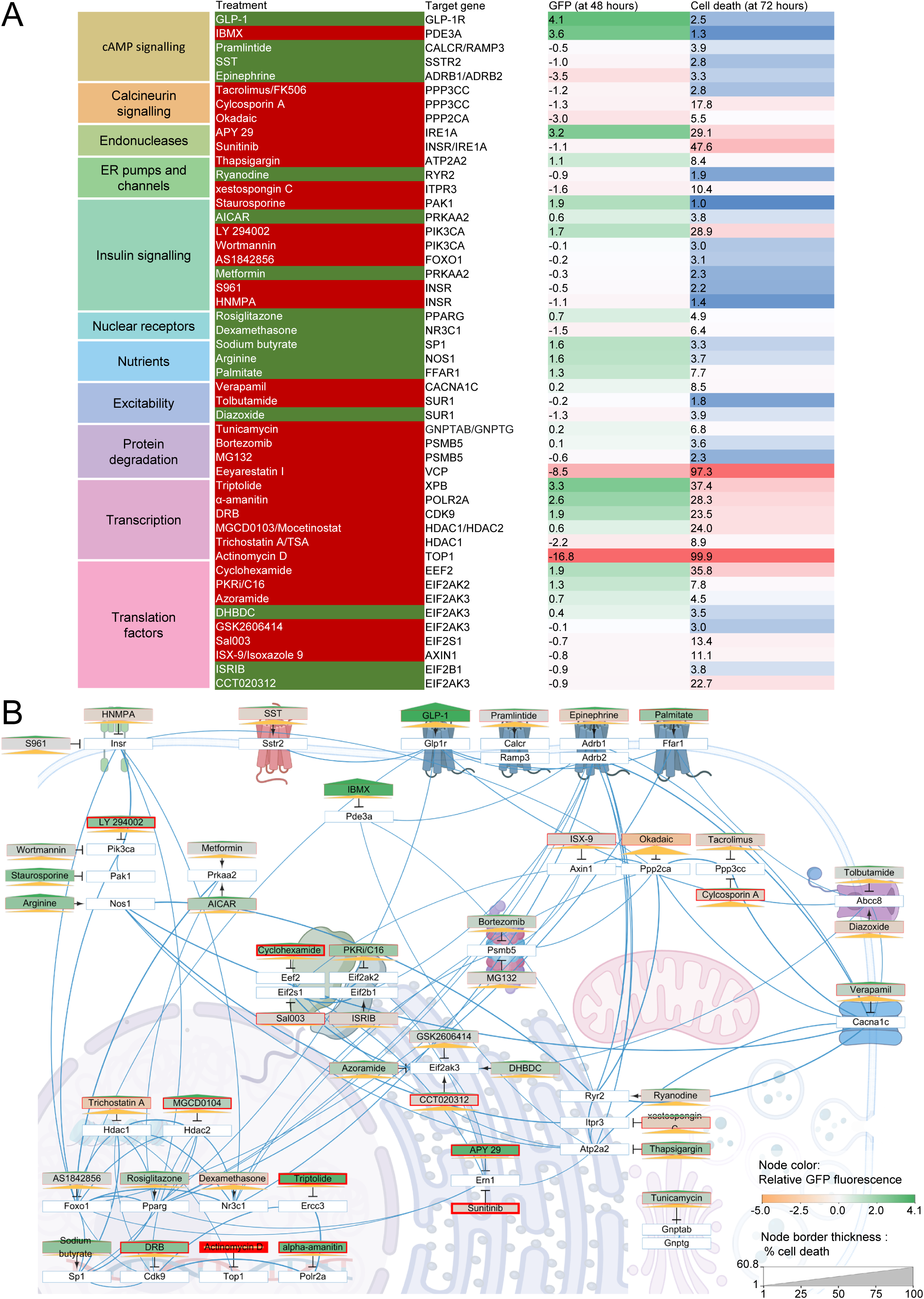
Integration of STRING protein networks and a small molecule screen on Cytoscape. **(A)** Primary binding partners for small molecules used in the screen localized in Cytoscape. Binding partners (bold) are chosen based on their affinity to the small molecule. Color of the nodes represents GFP fluorescence (scaled) after 48 hours of treatment. Borders of the nodes represents cell death (scaled PI fluorescence) after 72 hours of treatment. Height of green triangles represents *Ins2(*GFP)^LOW^ to *Ins2(*GFP)^HIGH^ transition rates. Height of orange triangles represents *Ins2(*GFP)^HIGH^ to *Ins2(*GFP)^LOW^ transition rates. Blue lines represent STRING interactions. Pointed arrows with grey lines represent upregulation. Inhibitor arrows with grey lines represent downregulation.

## Discussion

This study aimed to further our understanding of dynamic *Ins2* gene activity states, their proteomic profiles, and their regulation by key cellular processes. Several prior studies have labelled β cells using *Ins1*, but we chose the evolutionarily conserved *Ins2* gene due to the greater similarity between *Ins2* and *INS*^58,59^. Furthermore, our prior scRNAseq analysis revealed strong correlation between the two insulin genes in mice, and we had also observed similar cell states using an *Ins1* promoter reporter in another previous study^10,15^.

Our 3D imaging showed how the *Ins2*(GFP)^HIGH^ and *Ins2*(GFP)^LOW^ expression states existed in live intact islets. The two *Ins2* gene expression states were present in intact islets. We did not find any correlation for either cell population to the islet periphery or core. This is similar to what was observed by Johnston *et al*, in which ‘hub cells’, a β cell sub-set with similarities to *Ins2*(GFP)^LOW^ cells, did not show any clear preference for the islet center or periphery^60^. Interestingly, cells in the same activity states were more likely to be next to each other, forming loose patches of cells. Currently, it is unknown why these cells tend to cluster, and future experiments may further explore the spatial dynamics of the two cell states and their relationship with other cells^61,62^.

Previous studies identified the *Ins2*(GFP)^HIGH^ cells as being more mature, producing more *Ins2* mRNA, and more susceptible to stress^15^. The current study characterized the *Ins2*(GFP)^HIGH^ and *Ins2*(GFP)^LOW^ expression states at the proteome level and revealed mechanisms governing *Ins2* gene activity and cell state transitions. Proteomics analysis supported our prior hypothesis that *Ins2*(GFP)^HIGH^ are more mature, including the upregulation of the proliferation repressor RING2. Prior studies have shown that β cells which produce less insulin have increased rates of proliferation^19^. When adjusting for multiple comparisons, we found relatively few proteins that were significantly different in between our samples. This was attributed to the low cell counts in the *Ins2*(GFP)^HIGH^ state which resulted in high variability between those samples. The low cell counts may result from cell death during cell dispersion and FACS, perhaps worsened by the *Ins2*(GFP)^HIGH^ state’s innate fragility^15^. However, strong trends in the dataset prompted us to perform our analysis without adjusting for multiple comparisons. Minimally biased interrogations of proteins and protein networks selectively present in β cells and β cell states build on past efforts to map the β cell proteome. Previously proteomic analysis of FACS purifying mouse β cell populations based on insulin immunoreactivity yielded 6955 proteins across 36 islet samples with various diets and islet cell types^20^; this led to the identification of a subpopulation of β cells with low CD81 expression which had enrichment for factors in spliceosome and RNA processing pathways, but reduced oxidative phosphorylation factors, similar to our *Ins2*(GFP)^HIGH^ cells^20^. Two β cell subpopulations identified in a study on human β cell scRNAseq data had low *INS* expression, and while one high UPR marker expression, the other had enrichment in pathways related to oxidative stress^8^. These two β cell subpopulations possess similarities with our *Ins2*(GFP)^LOW^ cells, which had upregulation in ER stress markers and metabolic processes (Supplemental Fig. 8A-C), and it was proposed that these low *INS* expressing cells were in a state of recovery from the various stresses of high insulin production^8^. A recent study investigating younger and older insulin granules in INS-1 cells found proteins such as DYNC1H1, DCTN4, ARHGAP1 and EIF2S2 enriched in younger granules; these proteins were also significantly or trending towards enriched in *Ins2*(GFP)^LOW^ cells, suggesting that the two cell states may have different proportions of young and old insulin granules^63^. Overall, our study identifies novel and previously established β cell markers at the protein level and presented results showing how the cells in a high *Ins2* gene expression state had enrichment for factors related to RNA processing and transcription, with reduced oxidative phosphorylation and ER stress markers.

Our OPP protein translation assay revealed that the *Ins2*(GFP)^HIGH^ state had significantly higher protein synthesis activity. This upregulation of translation could be mediated by increased abundance of mRNA processing proteins (such as PAPOLA and ZRANB2) and proteins which may play a role in translation (such as DRG1 and ZC3H15) that were detected in proteomics (Supplemental Fig. 5B). Since our prior study showed that FACS purified *Ins2*(GFP)^HIGH^ cells have upregulated *Ins2* mRNA using qPCR^15^ and our scRNAseq data showed that *Ins2* mRNA was the most positively correlated gene with GFP mRNA^15^, this cell state is likely associated with both high insulin transcription and translation. Future studies should look into whether this upregulation in translational capacity is indicative of enhanced insulin secretion capacity^7,64,65^.

Our previous work revealed that cells *Ins2* gene activity was dynamic, and cells in the two cell states may transition between each other^15^. Our targeted perturbation analysis further explores this phenomenon and revealed pathways which impact endogenous *Ins2* gene activity and cell state transitions. A small molecule screen was previously done on live zebrafish larvae, with a set concentration for all treatments^66^. We did not identify overlap between their significant hits and ours, and we speculate this to be a result of the difference between whole organism screens and cell culture screens, as well as possible differences between the organisms involved. While whole organism screens can provide *in vivo* results and minimally perturb the health of biological samples during processing, cell culture screens allow direct interaction between treatments and the target cells. Our results revealed the calcineurin inhibitors tacrolimus, cyclosporine A, and okadaic acid reduced *Ins2* gene activity (Fig. 4C), consistent with known roles for calcineurin on insulin mRNA in transgenic mice with a human insulin promoter^67^. Okadaic acid and cyclosporine A are the two top inducers of *Ins2*(GFP)^HIGH^ to *Ins2*(GFP)^LOW^ transitions. NFAT, a calcineurin target, has been reported to be required for cAMP-mediated upregulation of insulin transcription in INS-1E cells^68^. Interestingly, IRS2, a reported target of calcineurin for regulation of human β cell survival, is enriched in *Ins2*(GFP)^HIGH^ cells, further supporting a role for calcineurin in cell state regulation^69^.

A key finding of our study was that GLP-1 amide and IBMX robustly increased *Ins2* activity over time, peaking at around 25 hours for GLP-1 amide and 55 hours for IBMX (Fig. 4B). This is consistent with previous data demonstrating that cAMP signaling plays a critical role in insulin production and secretion, with augmentation of cAMP having significant effects on β cell function and survival^70–74^. GLP-1 has also been shown to increase insulin mRNA and *Ins1* promoter activity in INS-1 insulinoma cell lines, and we now extend these results to the endogenous *Ins2* gene in mice primary cells^58^. Interestingly, GLP-1 and IBMX, also had the highest rates of *Ins2*(GFP)^LOW^ to *Ins2*(GFP)^HIGH^ transitions (Fig. 5D). Our proteomics analyses revealed that *Ins2*(GFP)^HIGH^ cells had upregulation in two phosphodiesterase (PDE) isoforms, PDE1C and PDE8A, both of which are known to play major roles in regulating cAMP and GSIS in rodent β cells^75,76^. Okadaic acid, the strongest inducer of *Ins2*(GFP)^HIGH^ to *Ins2*(GFP)^LOW^ transitions, is known to increase PDE expression in rat adipocytes^77^. It is also worth noting that despite strongly upregulating GFP expression, many treatments including APY29, triptolide, alpha amanitin, staurosporine, sodium butyrate, and arginine, did not induce nearly as many cell state transitions as IBMX and GLP-1 amide. Taken together, these results may suggest that the dynamics between calcineurin, cAMP, and PDE concentrations play a major role in regulating *Ins2* gene activity and cell state transitions.

Some small molecules had effects on GFP fluorescence that were strongly correlated to cell health. For example, actinomycin D and eeyarestatin I killed cells rapidly upon treatment, resulting in a rapid collapse of GFP fluorescence. Interestingly, a subset of drugs including APY29 (IRE-1A inhibitor), triptolide (TFIIH inhibitor), alpha amanitin (RNA polymerase II inhibitor), cyclohexamide (EEF2 inhibitor), DRB (CDK9 inhibitor), and LY294002 (PI3K inhibitor) induced a strong increase in GFP fluorescence followed by rapid cell death after prolonged treatment. The endonuclease IRE-1A is an important factor in β cell stress regulation^57^; inhibition of IRE-1A results in increased insulin mRNA content and β cell function, something that we also observe via the increase in GFP after 48 hours of inhibition via APY29^78^. The high cell death that follows after this peak in *Ins2* gene activity may be a result of β cell overwork^19^. Inhibition of general transcriptional machinery such as TFIIH and RNA polymerase II is highly detrimental to cell health after prolonged periods of time ^79,80^. Inhibition of translation and cell cycle factors are also known to be toxic to cells^81,82^. The PI3K signalling pathway is linked to cell survival and resistance mechanisms^83^. The toxicity of these drugs may result in a buildup of stress cumulating in eventual cell death It is interesting to note that the increase in GFP fluorescence in these drugs only occurs in *Ins2*(GFP)^HIGH^ cells, but not *Ins2*(GFP)^LOW^ cells. This may be due to the inherent fragility in *Ins2*(GFP)^HIGH^ cells, which may make them more vulnerable to the effects of stress^15^. Autofluoresence is also known to increase during the process of cell death, and this may also contribute to the increase in GFP fluorescence^84^. The OPP assay done in the context of several small molecules of interest also supports this, with cells treated to 24 hours of triptolide and APY 29 displaying reduced translational capacity prior to cell death and reduction in GFP fluorescence. Furthermore, these molecules all have the capacity to directly influence insulin production through the above stated mechanisms, with most having an inhibitory effect. This may suggest the existence of feedback mechanisms, as the cell tries to replenish insulin mRNA stores through increasing *Ins2* gene activity. These findings may suggest links between cell health and *Ins2* gene activity.

In summary, we provide data which further revealed the differences between the *Ins2*(GFP)^HIGH^ and *Ins2*(GFP)^LOW^ expression states beyond our previous study. Our live cell 3D imaging captured the two cell states in intact living islets, and our proteomic and translational assays characterizes them at the protein and functional level. The small molecule screen revealed pathways which may impact *Ins2* gene activity and cell state transitions, including prominent effects by effectors of cAMP signalling. Future studies may further explore the effects of cAMP and calcineurin effectors on *Ins2* gene dynamics and cell state transitions, such as dose response experiments to gauge the thresholds for transitions and the balance between β cell health and function. Studies may also be conducted to explore the cell states in the context of patients with diabetes; it has been shown that some patients with long-term type 1 diabetes still possess β cells which produce low levels of insulin, which may have similarities with our *Ins2*(GFP)^LOW^ cells^85^. Our study highlights the importance of cell expression states and cell state transitions in the islet, and how targeting factors which influence cell state transitions may help manage β cell function in diabetes.

### Limitations of study

Our study has many limitations, and we list three examples here. First, our 3D imaging experiments were unable capture the innermost cells in larger intact islets, so we focused mostly on smaller islets. Future experiments could use 2-photon microscopy to analyze islets of varying sizes. Second, as mentioned in the discussion, our proteomics analyses had high variability within sample groups. Futures studies should increase the number of biological replicates in proteomics experiments. Third, our small molecule screen started with known compounds with expected effects and is therefore not unbiased. Now that the methodology is established, future studies could examine larger libraries of compounds that cover chemical space in a more unbiased way. Despite these limitations, our study provides new information on primary β cell states and the pathways that control them.

### Author Contributions

CM. J. C. designed studies, performed experiments, analyzed/interpreted data, wrote manuscript.

B. S. performed experiments, analyzed/interpreted data, wrote parts of manuscript.

H. H. C. analyzed/interpreted data.

X. H. performed experiments.

W. G. S. performed experiments.

G. P. B. performed experiments and analyzed/interpreted data.

Y. H. X. analyzed/interpreted data.

J. R. performed experiments, analyzed/interpreted data, wrote parts of manuscript.

J. D. J. conceived the project, designed studies, interpreted data, and edited the manuscript.

J.D.J. is the guarantor of this work and, as such, had full access to all the data in the study and takes responsibility for the integrity of the data and the accuracy of the data analysis.

## Supporting information

Star methods

## Acknowledgments and funding

We thank many colleagues for helpful discussions during conferences, meetings, and presentations. Research was supported by a CIHR operating grant (PJT152999) to J.D.J., the JDRF Centre of Excellence at UBC, and core facilities in the Life Sciences Institute.

## Resource Availability

**Lead contact:** James D. Johnson, PhD., 5358 Life Sciences Building, 2350 Health Sciences Mall, University of British Columbia, Vancouver, Canada, V6T 1Z3 Phone: 604 822 7187, X/Bluesky: @JimJohnsonSci

## Materials availability statement

This study did not generate new unique reagents.

## Data and Code Availability

This study did not generate new code.

## Data

Proteomic data are available on ProteomExchange (number PXD052023).

## Code

Coding used in analyses are available upon reasonable request to the authors.

## Other items

Requests for other data are available upon reasonable request to the authors.

