## supplement for "Signal transduction pathways controlling *Ins2* gene activity and beta cell state transitions"

### Supplemental Figure 1

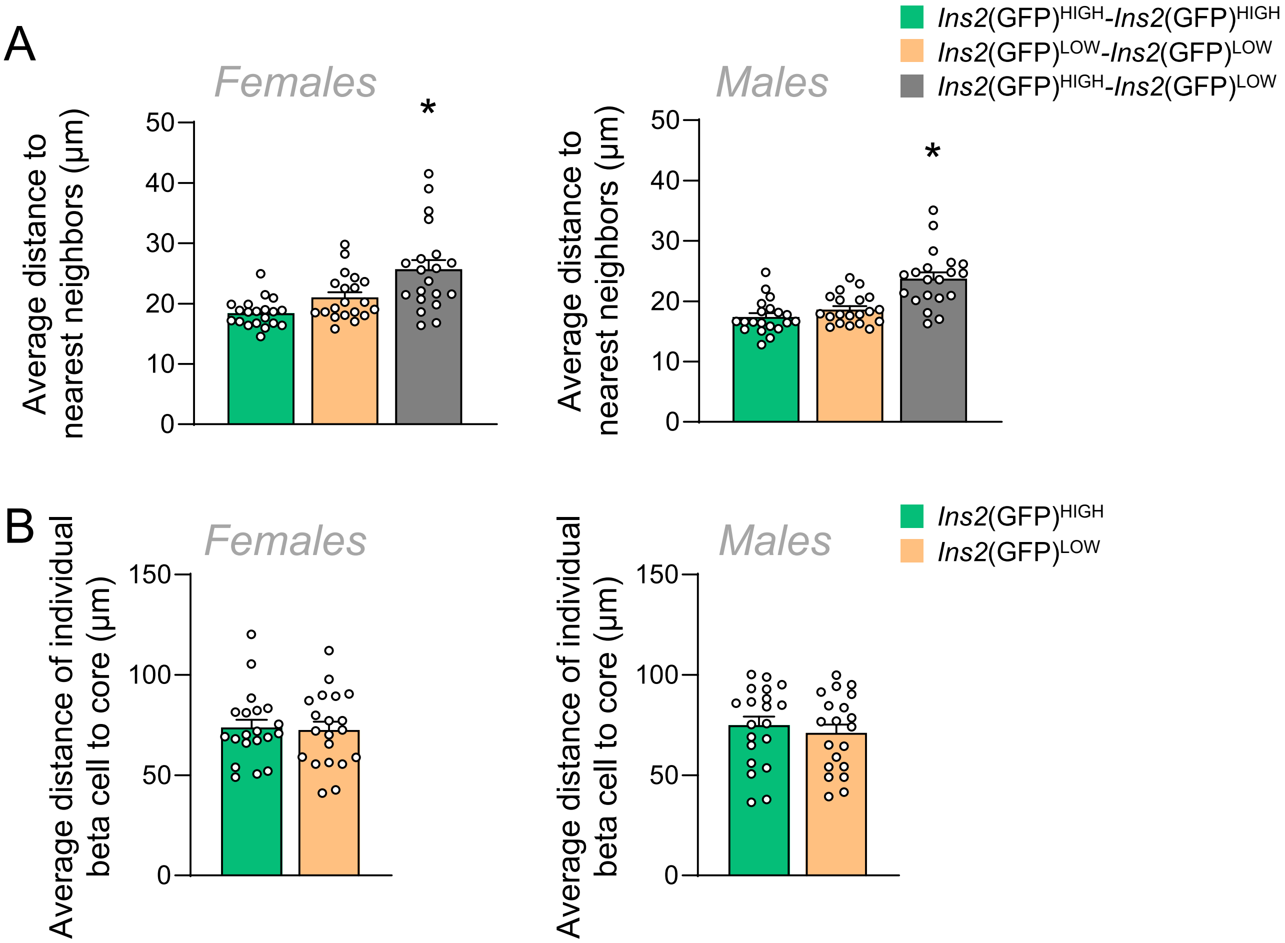


**Figure S1. Live cell 3D imaging of intact islets in the context of male and female *Ins2*$^{GFP}$ mice. (A)** Nearest neighbor analysis of cells from Ins2GFP knock-in mice (n=20 islets per sex, collected from three male and 3 female mice). One-way ANOVA. **(B)** Correlation to core analysis in the context of male and female mice. Student's t-test. Data are represented as mean +/- SEM. * $p < 0.05$.

Supplemental Figure 2

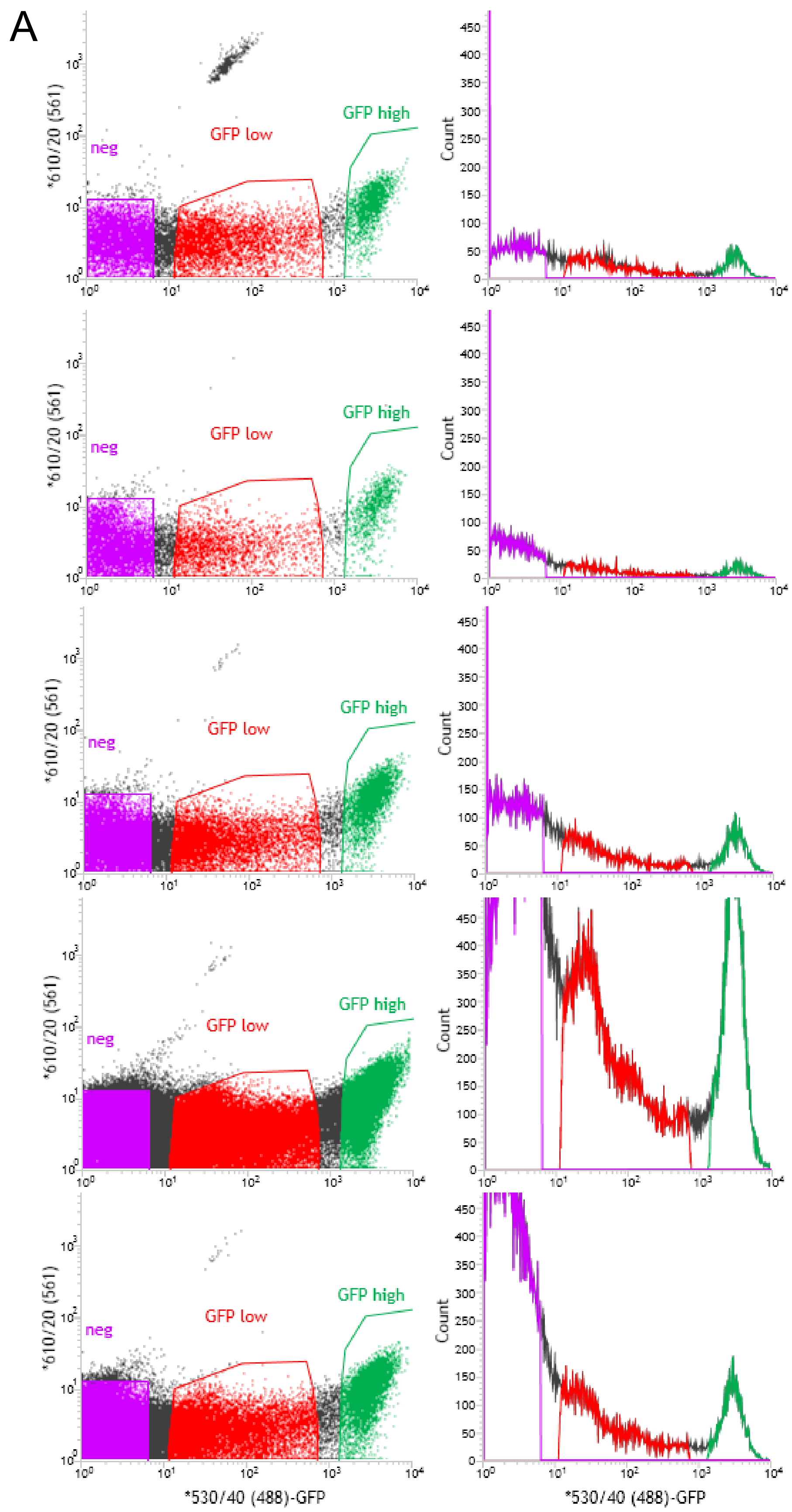


**Figure S2. FACS purified populations for proteomics. (A)** Dot plots and histograms of FACS purified cells that were used in the mass spec-based proteomics data. Samples were acquired from 3 male and 2 female mice.

### Supplemental Figure 3

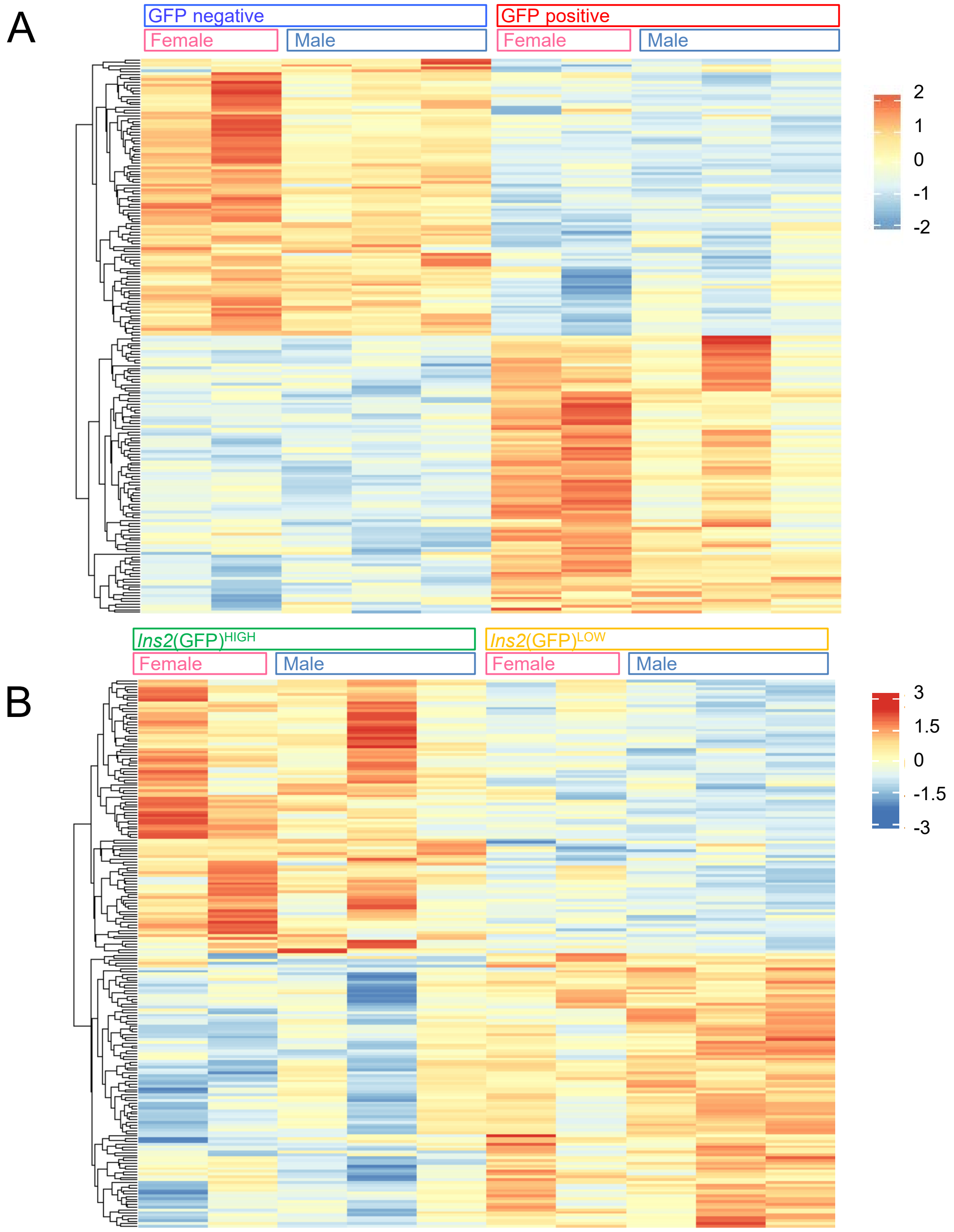


**Figure S3. Heatmaps of proteomics from all samples analyzed. (A)** Top 200 genes comparing GFP positive and GFP negative cells, ranked by –log10(p-value)*fold change. **(B)** Top 200 genes comparing *Ins2*(GFP)$^{HIGH}$ and *Ins2*(GFP)$^{LOW}$ cells, ranked by –log10(p-value)*fold change.

### Supplemental Figure 4

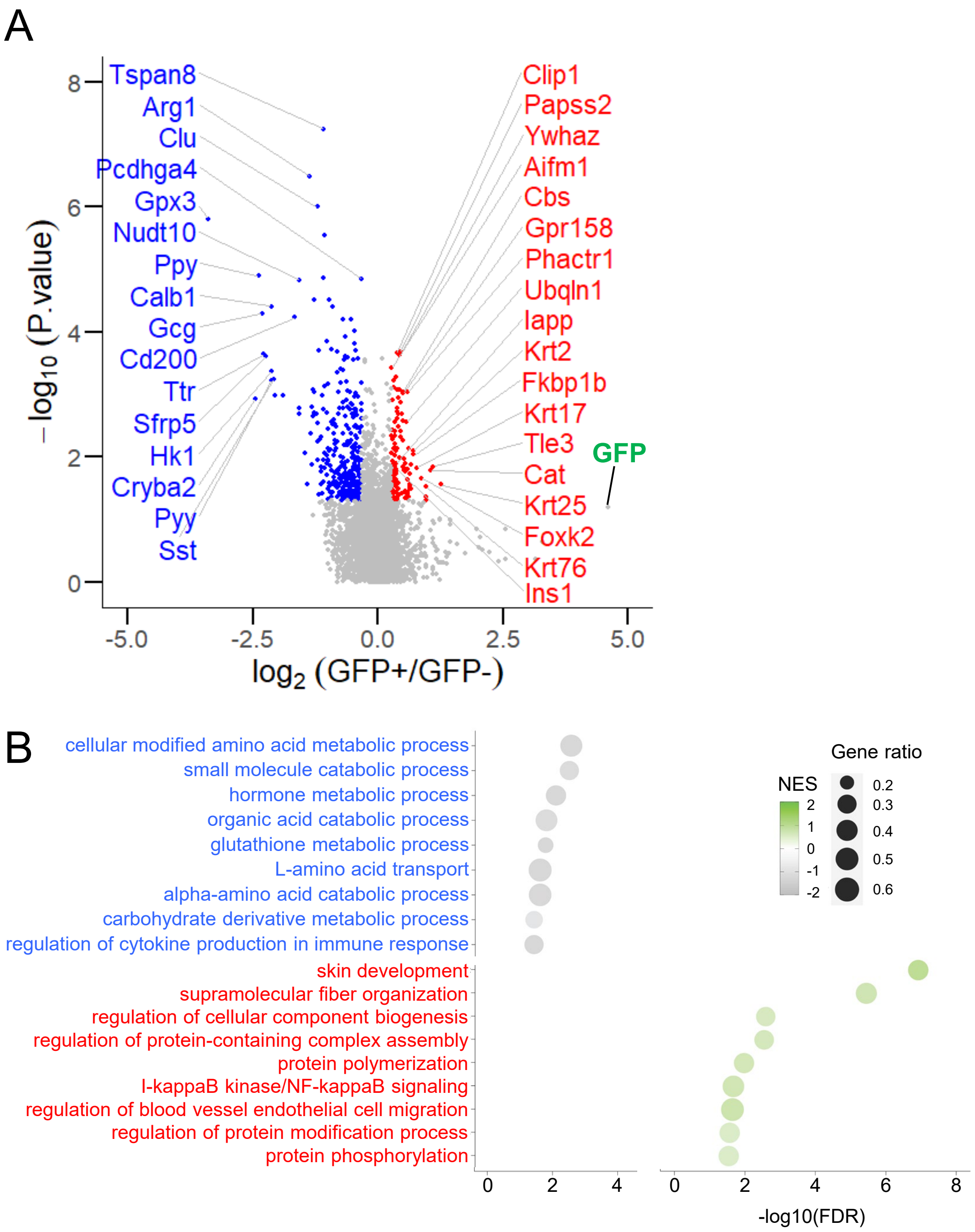


**Figure S4. Proteomic analysis of FACS purified *Ins2*$^{GFP}$ β cells. (A)** Volcano plot of differentially expressed proteins comparing FACS purified GFP positive and GFP negative cells. **(B)** Gene set enrichment analysis of pathways enriched in GFP positive and GFP negative cells.

Supplemental Figure 5

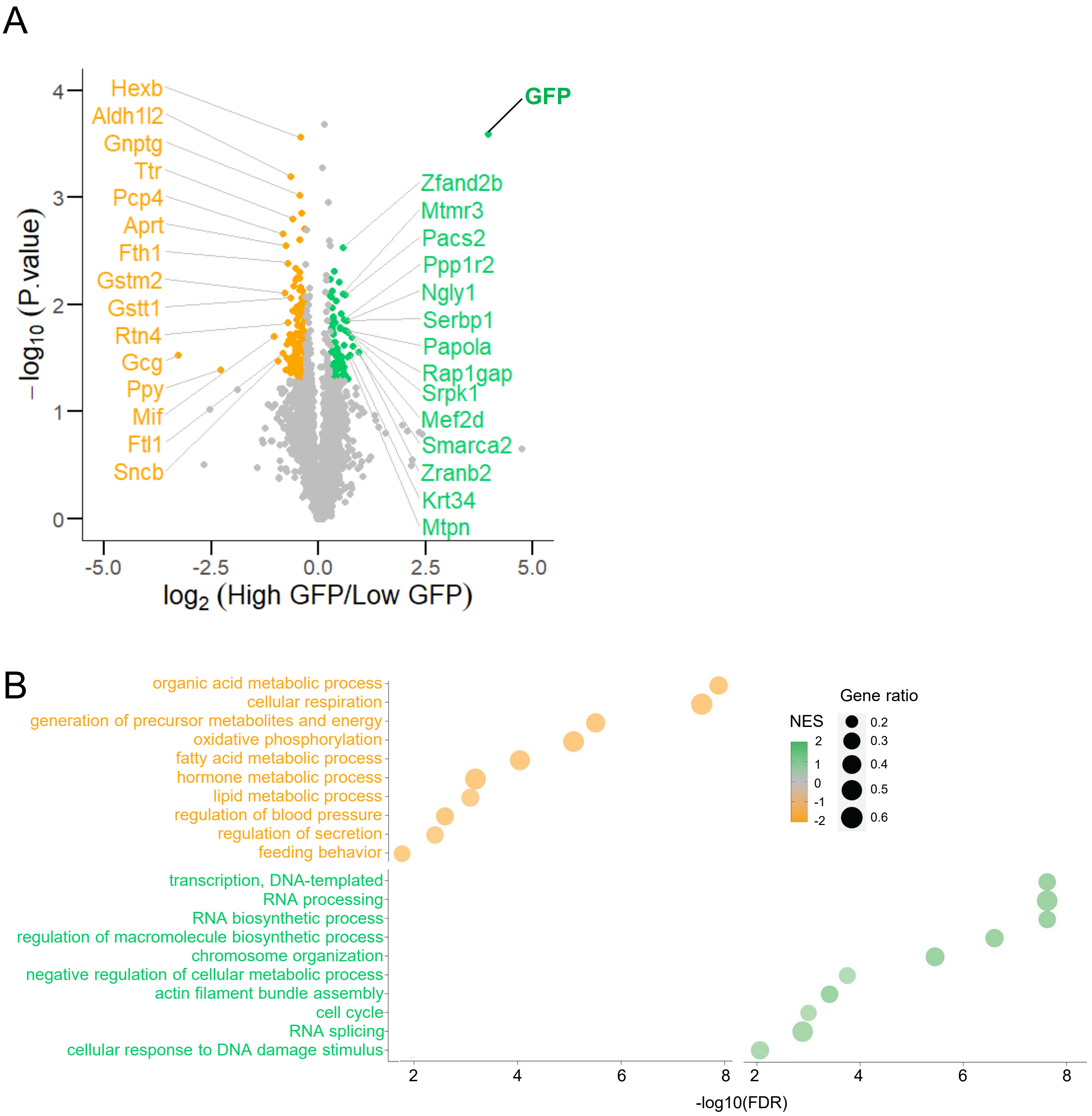


**Figure S5. Proteomic analysis of FACS purified *Ins2*$^{GFP}$ β cells. (A)** Volcano plot of differentially expressed proteins comparing FACS purified *Ins2*(GFP)$^{HIGH}$ and *Ins2*(GFP)$^{LOW}$ cells. **(B)** Gene set enrichment analysis of pathways enriched in *Ins2*(GFP)$^{HIGH}$ and *Ins2*(GFP)$^{LOW}$ cells.

### Supplemental Figure 6

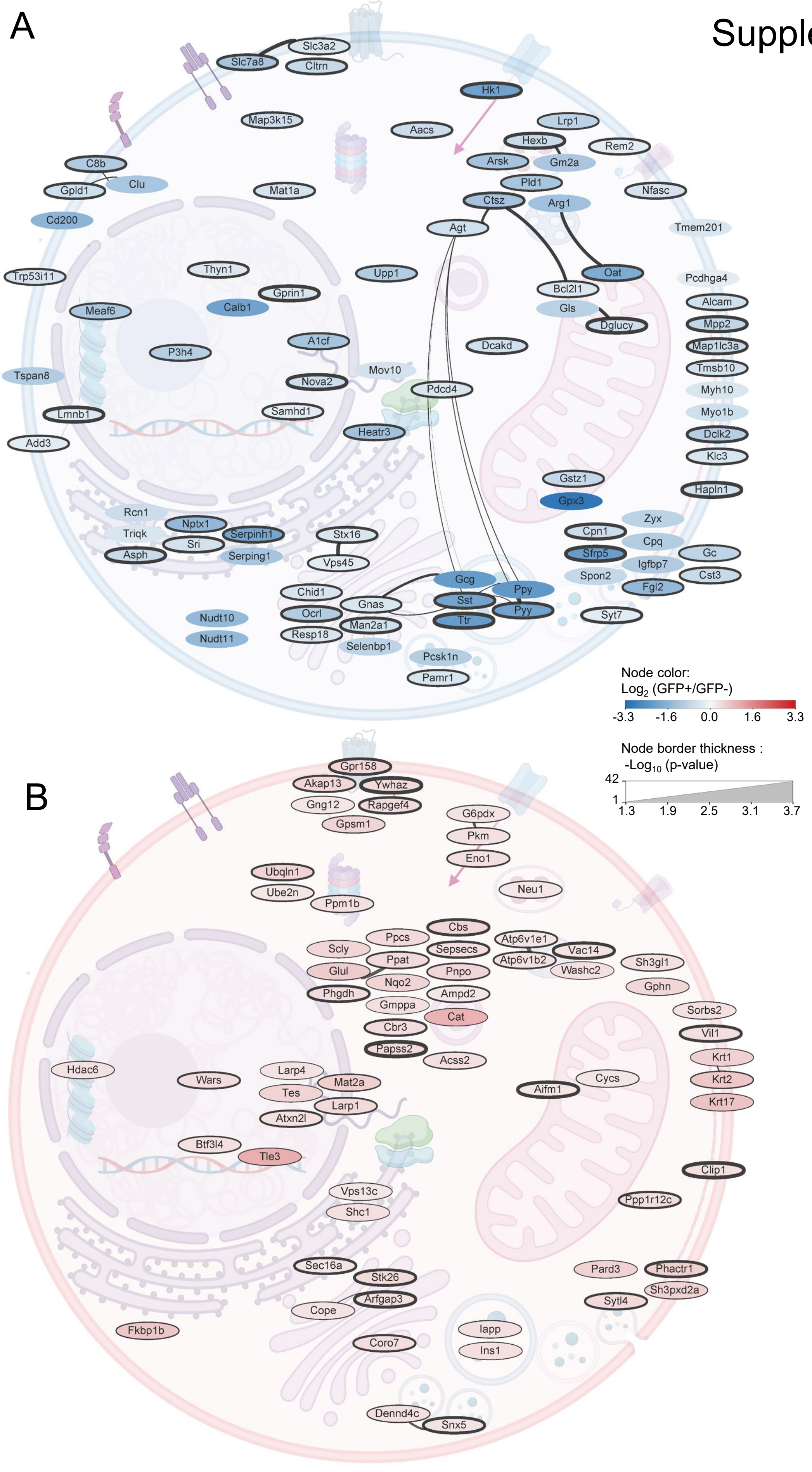

**Figure S6. Integration of STRING networks and protein expression data in Cytoscape reveals differences in protein production and export between GFP positive and GFP negative cells. (A)** Integration of STRING networks and protein expression data of differentially abundant proteins (top 100 with p value lower than 0.05) in GFP negative cells in Cytoscape. Deeper color of the nodes represents the fold change (GFP negative/GFP negative), while the thickness of the line around the node represents p value. **(B)** Integration of STRING networks and protein expression data of differentially abundant proteins (top 100 with p value lower than 0.05) in GFP positive cells in Cytoscape. Deeper color of the nodes represents the fold change (GFP positive/GFP negative), while the thickness of the line around the node represents p value.

### Supplemental Figure 7

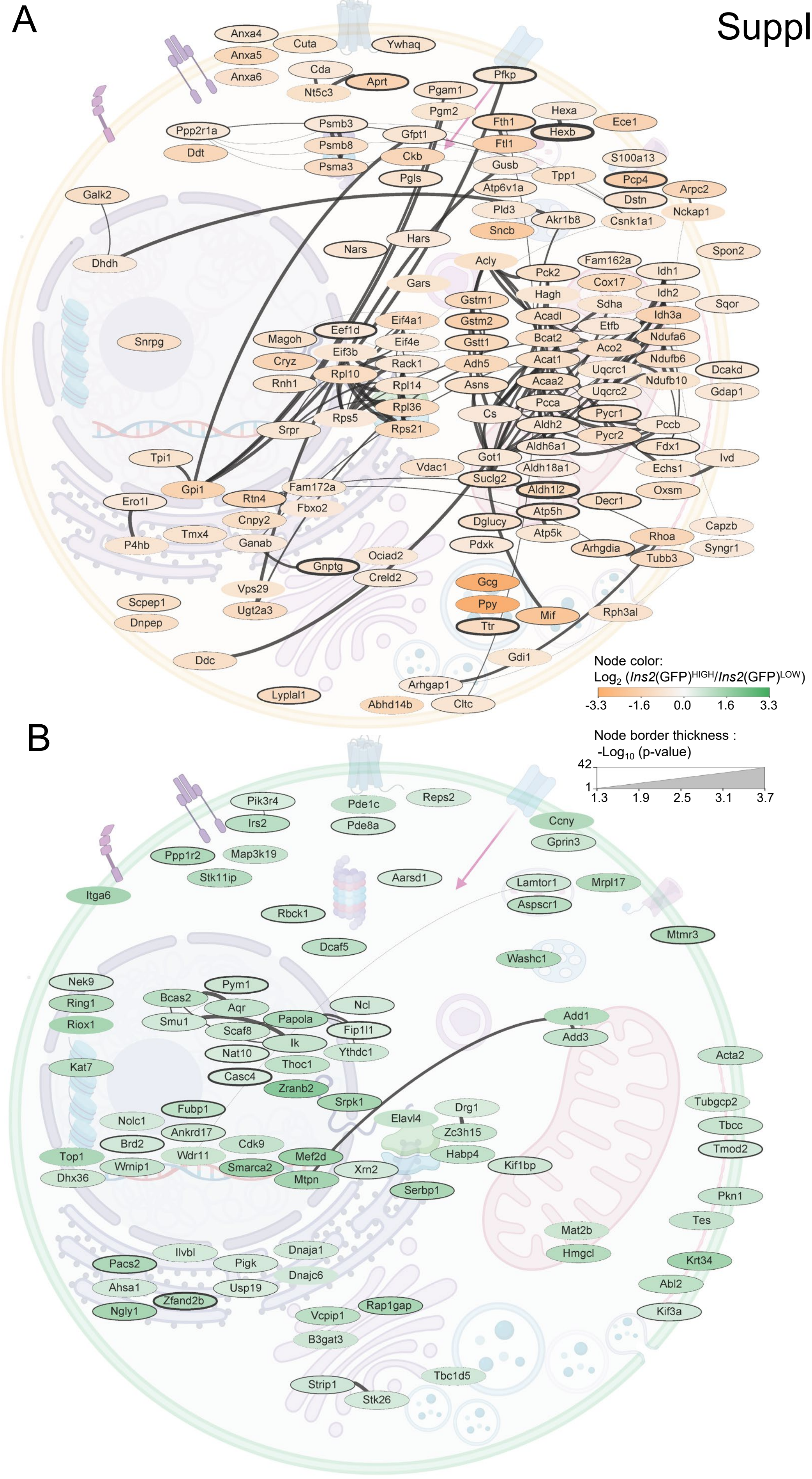

**Figure S7. Integration of STRING networks and protein expression data in Cytoscape reveals differences in transcription and mRNA processing between GFP expression states. (A)** Integration of STRING networks and protein expression data of differentially abundant proteins (p value lower than 0.05) in *Ins2*(GFP)$^{HIGH}$ cells in Cytoscape. Deeper color of the nodes represents the fold change (*Ins2*(GFP)$^{LOW}$/*Ins2*(GFP)$^{HIGH}$), while the thickness of the line around the node represents p value **(B)** Integration of STRING networks and protein expression data of differentially abundant proteins (p value lower than 0.05) in *Ins2*(GFP)$^{LOW}$ cells in Cytoscape. Deeper color of the nodes represents the fold change (*Ins2*(GFP)$^{HIGH}$/*Ins2*(GFP)$^{LOW}$), while the thickness of the line around the node represents p value.

### Supplemental Figure 8

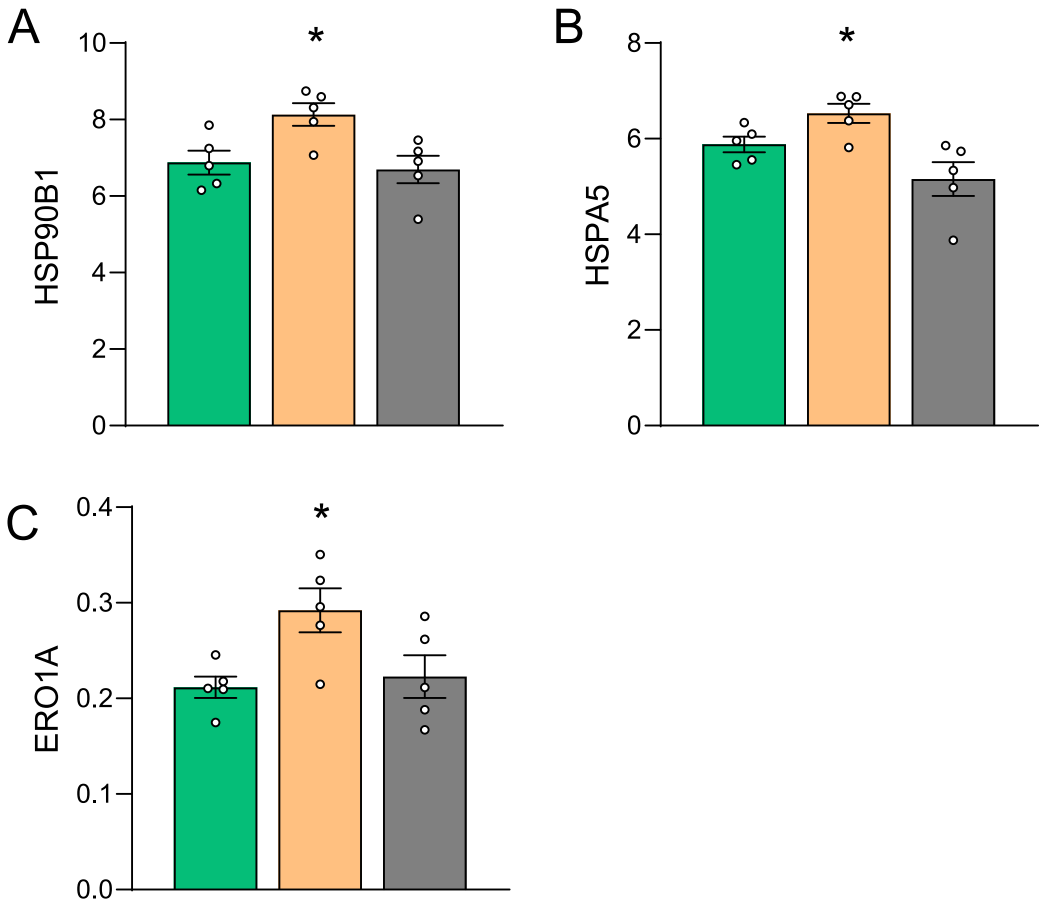


**Figure S8. Relative protein abundance of major ER markers in FACS purified populations. (A)** HSP90B1 abundance. **(B)** HSPA5 abundance. **(C)** ERO1A abundance. One-way ANOVA. Data are represented as mean +/- SEM. * $p < 0.05$.

Supplemental Figure 9

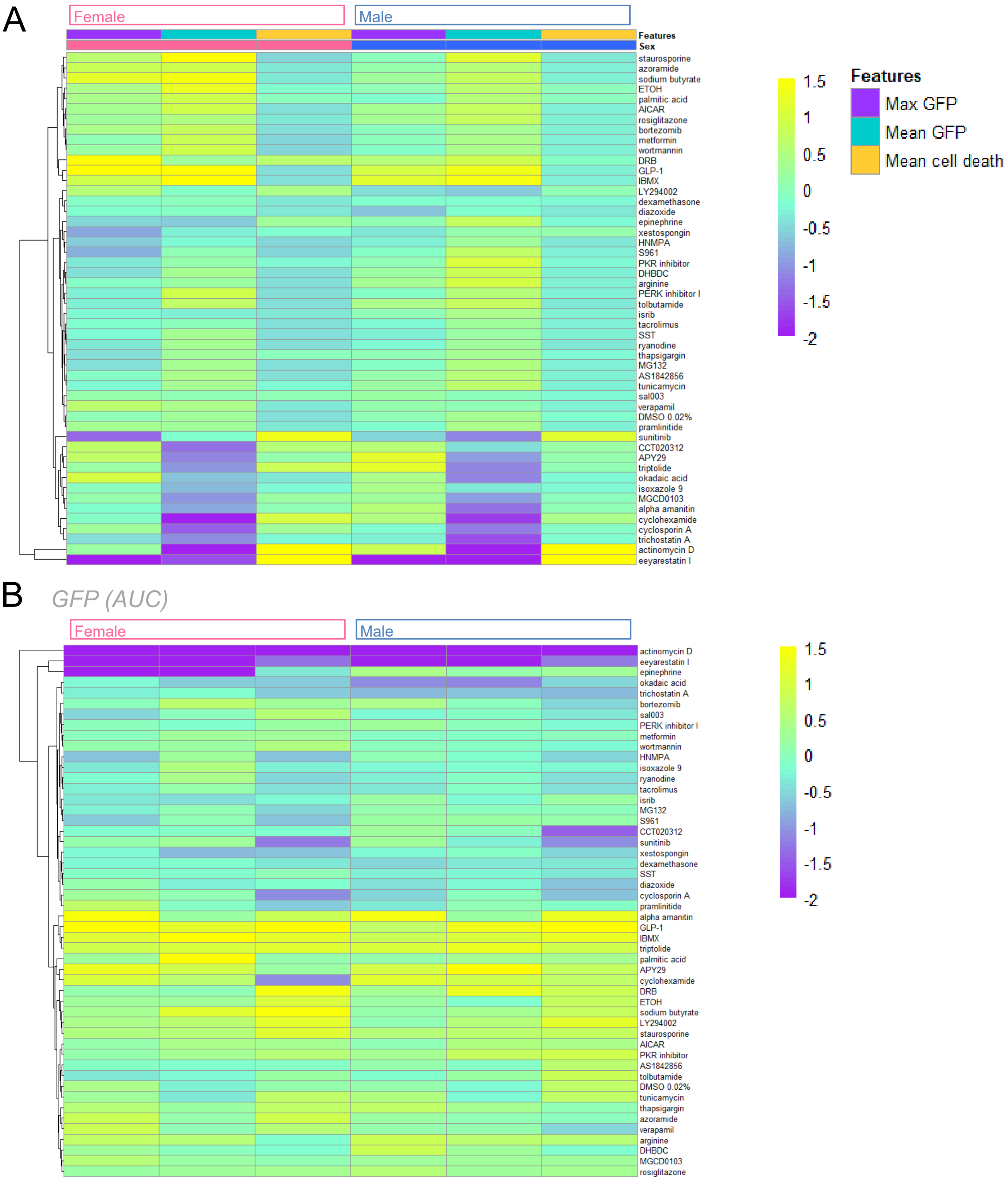


**Figure S9. Hierarchical clustering of cell behavior traits from small molecule screen. (A)** Hierarchical clustering of cell behavior traits in the context of sex. **(B)** Average GFP fluorescence for each treatment across all six individual mice (3 male and 3 female).

### Supplemental Figure 10

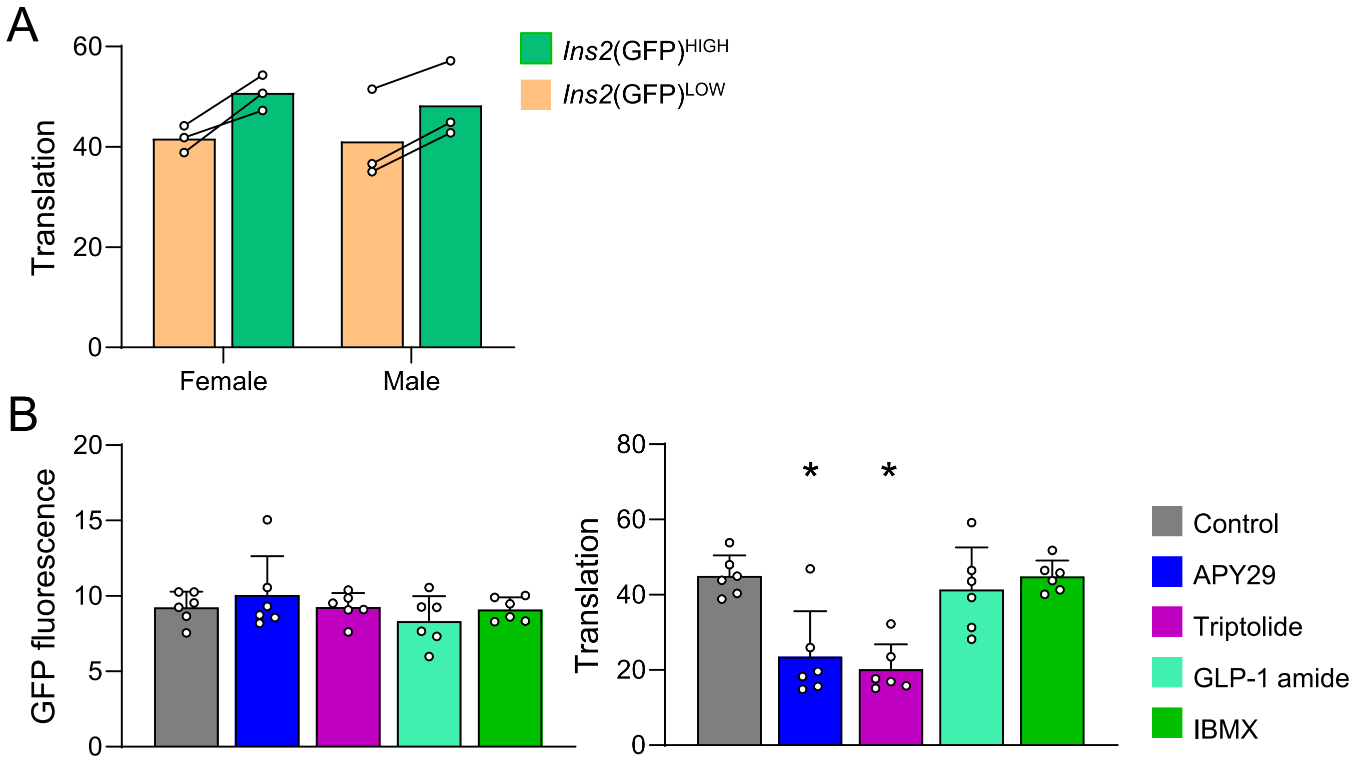


**Figure S10. OPP protein synthesis assay on small molecules of interest from the screen. (A)** Translational capacity in *Ins2*(GFP)$^{HIGH}$ and *Ins2*(GFP)$^{LOW}$ cells in the context of sex (3 male and 3 female mice). Student's t-test. **(B)** GFP fluorescence and translational capacity in cells treated for 24 hours by APY29, triptolide, GLP-1 amide, and IBMX. One way ANOVA. Data are represented as mean +/- SEM. * $p < 0.05$.
