## Supplementary material for "Signal transduction pathways controlling *Ins2* gene activity and beta cell state transitions": Star methods

### Animal husbandry and breeding

### Animals were housed in the Modified Barrier Facility at UBC, under protocols approved by the UBC Animal Care Committee in accordance with national and international guidelines. To maintain consistency with our prior study ^1^ and to keep the genotype as close to wild-type as possible, all mice in this study were *Ins2*^WT/GFP^ heterozygous mice. Including the *Ins1* gene, these mice have 3 out of 4 wildtype insulin alleles. Mice used for islet isolations were between 60 and 65 weeks of age.

### Islet isolation and dissociation

### Pancreatic islets were isolated using collagenase when mice were aged 60 weeks, filtered, and hand-picked as previously described ^2^. Islets were cultured overnight (37°C, 5% CO2) in RPMI1640 medium (ThermoFisher, Waltham, United States) with 11 mM glucose (Sigma), 100 units/ml penicillin, 100 μg/ml streptomycin (ThermoFisher, Waltham, United States), and 10% vol/vol fetal bovine serum (FBS; ThermoFisher, Waltham, United States). For islet dissociation, islets were washed 4 times with minimum essential medium (Corning, Corning, United States) and immersed for 5 minutes in 0.05% trypsin at 37°C with pipetting. Cells were then resuspended in RPMI1640 medium, plated in appropriate reservoirs or plates, and rested for 24 hours (37°C, 5% CO2). Cells were incubated for 2 hours in the presence of 0.05 μg/mL Hoechst 33342 (Sigma–Aldrich, St. Louis, United States) to mark nuclei and 0.5 μg/mL propidium iodide (Sigma–Aldrich, St. Louis, United States) to mark cell death. Drug/small molecules for the perturbation study were purchased from the sources indicated in Table 1 and applied immediately prior to imaging.

### Fluorescence-activated cell sorting

### Pancreatic islets were dispersed using 0.05% trypsin and resuspended in 1x PBS with 0.005% FBS. Dispersed islets were then filtered into 5 ml polypropylene tubes. Sorting was conducted on a Cytopeia Influx sorter (Becton Dickinson, Franklin Lakes, USA) at the Life Sciences Institute FLOW core facility. Cells were excited with a 488 nm laser (530/40 emission) and a 561 nm laser (610/20 emission).

### Live cell imaging and analysis

### For 3D live imaging, intact mouse islets were incubated overnight in culture media (RPMI1640 medium with 11 mM glucose, 100 units/ml penicillin, 100 μg/ml streptomycin, and 10% vol/vol FBS) with Hoechst 33342 (0.05 μg/mL) and propidium iodide (0.5 μg/mL). After 24 hours, cells were transferred into 2 well or 8 well µ-Slides (Ibidi) with fresh culture media. Images were then acquired using a LEICA THUNDER Imager Live Cell & 3D assay microscope at 25x water objective (numerical aperture 0.95). Analysis of 3D live cell imaging experiments was done using Imaris Microscopy Image Analysis software with the FIJI Labkit module and custom R scripts. Image processing, including smoothing and background subtraction, was performed using Imaris. Labkit facilitated image labeling and pixel classification for the identification of GFP fluorescence in individual β cells. Nearest neighbor analysis was done using Imaris Python XTensions. Distance-to-core analysis was conducted on Imaris, using a reference frame positioned at the islet's center to measure the distance of each cell from the origin reference frame. Plots were generated for visualization using Graphpad Prism v10 or custom R scripts.

To investigate the effects of small molecules on GFP fluorescence, we cultured dispersed islet cells from *Ins2*^GFP^ mice on 384-well glass bottom plates (Perkin Elmer, Waltham, United States) and performed live cell imaging using an ImageXpress^Micro^ environmentally controlled, robotic system (Molecular Devices, San Jose, United States) which employed a 300 W Xenon lamp as previously described ^1^. Cells were stained with Hoechst 33342 and propidium iodide as described above. Images were acquired at 10x air objective, numerical aperture 0.3, at 30-minute intervals up to 96 hours, with cells being exposed to 359 nm light for 110 ms, 491 nm light for 15 ms, 561 nm for 75 ms. Analysis of live cell imaging experiments on *Ins2*^GFP^ cells was done using MetaXpress analysis software and custom R scripts. Tracking of GFP fluorescence in individual cells was performed as previously described ^1^. The *Ins2*(GFP)^HIGH^ and *Ins2*(GFP)^LOW^ subpopulations were identified using model-based clustering ^3^.

### Protein synthesis assay

### One day after isolation, dispersed islets were seeded into an optical 96-well plate (Perkin Elmer, Waltham, United States) at a density of approximately 16,000 cells per well in culture media (RPMI1640 medium with 11 mM glucose, 100 units/ml penicillin, 100 μg/ml streptomycin, and 10% vol/vol FBS). Treatments were applied 3 hours after seeding. After 24 hours of incubation, fresh culture media was applied, then supplemented with 20 μM OPP (Invitrogen, Waltham, United States). The assay was performed according to instructions provided by the manufacturer. Cells were imaged at 10x, numerical aperture 0.3, with an ImageXpress^Micro^ high-content imager and analyzed with MetaXpress to quantify the integrated staining intensity of OPP-Alexa Fluor 594 in cells identified by NuclearMask Blue Stain.

### Mass Spectrometry / Proteomics

### For proteomics sample preparation, between 2,000 and 10,000 cells were diluted with ultrapure water, and then lysed by adding cold trifluoroethanol (TFE) (final 50% TFE) to the suspension. Samples were then chilled on ice for 10 min, vortexed for 1 min and further sonicated for 5-10 min in an ice bath. Tris (pH 8-8.5) was added to adjust the pH and concentration to final 100 mM Tris at pH 8-8.5. Total protein mass was estimated to be 0.3 ng/cell. Samples were reduced with tris(2-carboxyethyl)phosphine (1 μg TCEP: 50 μg protein) incubated for 20 min at room temperature, then alkalized with chloroacetamide (5 μg CAA:50 μg protein) incubated at 95 °C for 10 min. Then samples were diluted with 50mM Ammonium bicarbonate (final TFE < 10%). Proteins were digested with LysC/Trypsin (1 μg LysC/Trypsin: 50 μg protein) for 2 hours at 37 °C, followed by digestion with trypsin (1 μg trypsin: 50 μg protein) and incubated overnight at 37 °C. Samples were further digested with trypsin (1 μg trypsin: 125 μg protein) for 5 hours at 37 °C. Digestion was stopped by acidification. Samples were desalted with 6 mm depth of C18; briefly, STAGE (stop and go extraction) tips were conditioned with 100% methanol, equilibrated with 0.2% TFA, loaded with samples, then washed with 0.2% TFA, and finally eluted with 80 μL *2 of 40% ACN, 0.1% formic acid. Eluate was dried in vacuum then kept in -20°C. Before instrumental analysis, samples were reconstituted in in 0.5% acetonitrile, 0.1% formic acid and concentration was measured using NanoDrop One (ThermoFisher, Waltham, USA) with the A205 Scopes method (absorbance at 205 nm, baseline correction at 340 nm).

### For liquid chromatography steps, 75 ng of peptides were injected and separated on-line using NanoElute UHPLC system (Bruker Daltonics Billerica, United States) with Aurora Series Gen2 (CSI) analytical column, (25 cm x 75 μm 1.6μm FSC (fuse silica core) C18, with Gen2 nanoZero and CSI (captive spray interface) fitting; Ion Opticks, Parkville, Australia) heated to 50 °C and coupled to timsTOF Pro (Bruker Daltonics, Billerica, USA). A standard 30 min gradient was run from 2% buffer B to 12% buffer B over 15 min, then to 33% B from 15 to 30 min, then to 95% B over 0.5 min, and held at 95% B for 7.72 min. Before each run, the analytical column was conditioned with 4 column volumes of buffer A. Where buffer A consisted of 0.1% aqueous formic acid and 0.5 % acetonitrile in water, and buffer B consisted of 0.1% formic acid in 99.4% acetonitrile. The NanoElute thermostat temperature was set at 7 °C. The analysis was performed at 0.3 μL/min flow rate.

### For mass spectrometry steps, the Trapped Ion Mobility – Time of Flight Mass Spectrometer (TimsTOF Pro; Bruker Daltonics, Billerica, USA) was set to Parallel Accumulation-Serial Fragmentation (PASEF) scan mode for DIA acquisition scanning 100 – 1700 m/z. The capillary voltage was set to 1800V, drying gas to 3 L/min, and drying temperature to 180°C. The MS1 scan was followed by 17 consecutive PASEF ramps containing 22 non-overlapping 35 m/z isolation windows, covering the m/z range 319.5 – 1089.5 (more information in DIA windows). As for TIMS setting, ion mobility range (1/k0) was set to 0.70 – 1.35 V·s/cm^2^, 100 ms ramp time and accumulation time (100% duty cycle), and ramp rate of 9.42 Hz; this resulted in 1.91s of total cycle time. The collision energy was ramped linearly as a function of mobility from 27 eV at 1/k0 = 0.7 V·s/cm^2^ to 55 eV at 1/k0 = 1.35 V·s/cm^2^. Mass accuracy: error of mass measurement is typically within 3 ppm and is not allowed to exceed 7 ppm. TimsTOF Pro was run with timsControl v. 4.1.12 (Bruker, Billerica, USA). LC and MS were controlled with HyStar 6.0 (6.2.1.13, Bruker, Billerica, United States). For the final search and quantification of the results, acquired diaPASEF data were then searched using DIA-NN (v. 1.8.1) to obtain DIA quantification, with the spectral library generated from the canonical proteome for *Mus Musculus* from UniProt. Quantification mode was set to “Any LC (high precision) and a two-pass search was completed. All other settings were left default. Mice with a GFP protein ratio of less than 3 when comparing their respective *Ins2*(GFP)^HIGH^ and *Ins2*(GFP)^LOW^ samples were excluded.

### Statistics and data visualization

### Statistics and data representation for nearest neighbor analyses, distance to core analyses, and OPP translation assays, were conducted using GraphPad Prism v10 (GraphPad Software, San Diego, USA). Statistics and data representation for live cell imaging experiments employed custom R scripts. Student’s t-test and one-way ANOVAs were used for parametric data as indicated in figure legends. For all statistical analyses, differences were considered significant if the p value was less than 0.05. Error bars were presented as ± standard error of the mean (SEM).

### For proteomics analyses, proteins that were present in less than 80% of samples were removed from analysis. Significantly differential proteins were defined with a threshold of p value < 0.05 and a fold-change of more than 20%, and graphs were generated using custom R scripts. For STRING and Cytoscape analyses, significantly differentially abundant proteins were entered into the STRING website, where nodes and networks were generated. For STRING settings, only ‘Experiments’ and ‘Databases’ interactions were included, with medium confidence interaction scores. STRING networks and protein expression data were then imported into Cytoscape and reordered according to their cellular location. Background images were generated with Biorender.

### For STRING and Cytoscape analyses in our small molecule screen, the primary binding proteins for small molecules were determined using online literature search. Proteins were then entered into STRING, where nodes and networks were rendered. For settings on STRING, all interactions were included, with medium confidence interaction scores. STRING networks, GFP fluorescence, cell death, and cell state transition data were then imported and finalized in Cytoscape. Background graphs were generated using Biorender.

**Methods References**

1.	Chu, C.M.J., Modi, H., Ellis, C., Krentz, N.A.J., Skovso, S., Zhao, Y.B., Cen, H., Noursadeghi, N., Panzhinskiy, E., Hu, X., et al. (2022). Dynamic Ins2 Gene Activity Defines beta-Cell Maturity States. Diabetes *71*, 2612-2631. 10.2337/db21-1065.

2.	Brownrigg, G.P., Xia, Y.H., Chu, C.M.J., Wang, S., Chao, C., Zhang, J.A., Skovso, S., Panzhinskiy, E., Hu, X., Johnson, J.D., and Rideout, E.J. (2023). Sex differences in islet stress responses support female beta cell resilience. Mol Metab *69*, 101678. 10.1016/j.molmet.2023.101678.

3.	Scrucca, L., Fop, M., Murphy, T.B., and Raftery, A.E. (2016). mclust 5: Clustering, Classification and Density Estimation Using Gaussian Finite Mixture Models. R J *8*, 289-317.
